# Interplay between Nrf2 and αB-crystallin in the lens and heart of zebrafish under proteostatic stress

**DOI:** 10.1101/2023.04.04.535454

**Authors:** Jinhee Park, Samantha MacGavin, Laurie Niederbrach, Hassane S. Mchaourab

**Author notes:** To whom correspondence should be addressed: Hassane Mchaourab: Department of Molecular Physiology & Biophysics, Vanderbilt University, Nashville TN 37232.

## Abstract

A coordinated oxidative stress response, partly triggered by the transcription factor Nrf2, protects cells from the continual production of reactive oxygen species. Left unbuffered, reactive oxygen species can lead to protein aggregation which has been implicated in a spectrum of diseases including cataract of the ocular lens and myopathy of the heart. While proteostasis is maintained by diverse families of heat shock proteins, the interplay between the oxidative and proteostatic stress responses in the lens and the heart have not been investigated. Capitalizing on multiple zebrafish lines that have compromised function of Nrf2 and/or the two zebrafish small heat-shock proteins αBa- and αBb-crystallin, we uncovered a transcriptional relationship that leads to a substantial increase in αBb-crystallin transcripts in the heart in response to compromised function of Nrf2. In the lens, the concomitant loss of function of Nrf2 and αBa-crystallin leads to upregulation of the cholesterol biosynthesis pathway thus mitigating the phenotypic consequences of the αBa-crystallin knockout. In contrast, abrogation of Nrf2 function accentuates the penetrance of a heart edema phenotype characteristic of embryos of αB-crystallin knockout lines. Multiple molecular pathways, such as genes involved in extracellular interactions and implicated in cardiomyopathy, are revealed from transcriptome profiling thus identifying novel targets for further investigation. Together our transcriptome/phenotypic analysis establishes an intersection between the oxidative stress and chaperone responses in the lens and the heart.

## Introduction

Oxidative stress presents a sustained challenge to long lived cells, including lens fiber cells, and cells with high metabolic requirement, such as cardiac myocytes [1, 2]. Unbuffered reactive oxygen species (ROS) is associated with multiple deleterious effects on cellular homeostasis including DNA damage, protein oxidation and organelle dysfunction [3]. A central player in the maintenance of the cellular oxidative balance is the transcription factor Nrf2 [4]. Nrf2 regulates antioxidant responses by turning on the expression of enzymes that eliminate ROS and thus maintain the redox balance [4]. Through modulation of its interaction with Kelch-like ECH-associated protein 1 (Keap1), oxidative stress drives Nrf2 translocation into the nucleus to activate genes involved in glutathione synthesis, detoxification, elimination of ROS, and drug excretion [5].

In long lived cells, protein damage that leads to loss of stability and/or solubility can induce the formation of protein aggregates. A prominent example of the deleterious effects of protein aggregation occurs in the ocular lens. Accumulation of age-dependent damage to lens proteins induced by multiple factors, including UV-radiation and oxidation, leads to changes in their stability and solubility [6–8]. Gradual nucleation of aggregation-prone proteins eventually leads to large condensates that result in lens opacification, light scattering, and potentially age-related cataract [9]. Age-related cataract is one of the world’s most common causes of blindness [10–15]. Numerous lines of evidence suggest oxidative stress is a leading risk factor for cataract formation[1, 16–18]. Remarkably, lens protein’s relatively rich cysteine and methionine content makes it highly prone to oxidation by reactive oxygen species (ROS) [19–21]. Over 90% of cysteine residues and 50% of methionine residues are oxidized in cataract patient lenses [22]. Balancing the life-long danger of oxidative stress, the lens possesses a robust antioxidant defense system to scavenge and detoxify ROS. Glutathione (GSH) is present at exceedingly high concentrations in the lens, allowing it to resist oxidative damage [23]. The implied mechanistic link between Nrf2 activity and cataract formation has sparked interest in Nrf2 as a potential therapeutic target for cataract treatment and prevention [24–26].

The programmed elimination of light scattering organelles in lens fiber cells are essential to achieve the lens’s optical properties [27]. As a result, proteome turnover is exceedingly low, and lens proteins must remain for a lifetime. Proteostasis in the quiescent vertebrate lens is maintained almost exclusively by chaperone activity of the resident small heat shock proteins (sHSPs), αA- and αB-crystallin, which have been hypothesized to play a central role in preventing the aggregation of unstable and damaged proteins [28–31]. In addition, mutations in lens α-crystallins have been associated with congenital cataracts [32, 33]. Thus, through their complementary roles in preventing protein damage and inhibiting protein aggregation, Nrf2 and sHSPs are important factors in the maintenance of the lens optical properties.

Unlike αA-crystallin, which is mainly expressed in the ocular lens, αB-crystallin is detected in multiple tissues, including the heart, brain, skeletal muscle, kidney, and extracellular matrix [34]. Transcriptional regulation of *αB-crystallin* in mammals is tissue-specific and triggered by heat shock and other stress stimuli, including arsenite/cadmium, hypertonic/osmotic stresses, and oxidative stress, and is regulated by stress-activated proteins such as heat-shock factor1(hsf1) or transcription factor AP1 [35]. αB-crystallin has been implicated in maintaining cardiac homeostasis through a spectrum of roles in myocytes. It interacts with cytoskeletal proteins such as Titin, which serves as a molecular spring for passive muscle elasticity [36]. Indeed, it has been reported that high cardiomyocyte stiffness, which is highly correlated with aortic stenosis and dilated cardiomyopathy, was corrected by αB-crystallin through suppression of Titin aggregation [37]. In addition, αB-crystallin in myocytes has a protective role in response to ischemia-reperfusion stress, a condition that elevates reactive oxygen species (ROS) production [38]. Finally, the R120G mutation in the *αB-crystallin* gene is associated with congenital cataracts and muscular diseases, including cardiomyopathy [39, 40]. R120G-related cardiomyopathy is characterized by reductive stress, desmin aggregation in inclusion bodies, and ventricular dysfunction [41]. Collectively, these studies suggest that αB-crystallin plays a pleiotropic role in various cellular activities such as stabilization of the cytoskeletal structure, protein quality control, cell differentiation, and apoptosis. Thus, similar to lens fiber cells, sHSPs and Nrf2 are involved in the oxidative and proteostasis response in cardiomyocytes.

While multiple lines of evidence suggest a convergence between the roles of sHSPs, particularly αB-crystallin, and the oxidative stress response [38, 42–44], there has been no systematic investigation of the direct link between them nor has the *in vivo* molecular mechanism underlying this link been explored. In this study, we interrogated the mechanistic relationship between tissue-specific regulation of zebrafish *αBa*- and *αBb-crystallin* and oxidative stress due to Nrf2 depletion, focusing on the lens and heart tissues where αB-crystallin’s physiological roles has been demonstrated unequivocally. In zebrafish, two *αB-crystallin* paralogs have been identified due to gene duplication [45], αBa-crystallin and αBb-crystallin. Zebrafish *αBa-crystallin* transcripts are predominantly expressed in the lens, whereas *αBb-crystallin* is more widely expressed in multiple tissues, including the lens, muscle, and brain [45]. Importantly, recombinant zebrafish αBa-crystallin exhibits more potent chaperone-like activity than αBb-crystallin *in vitro* [45, 46]. These findings suggest that the two zebrafish αB-crystallins are subjected to divergent selection pressures to meet the challenges of distinct physiological functions.

Taking advantage of several zebrafish lines that have compromised αB-crystallins and/or Nrf2 function, we uncovered a transcriptional coupling between *nrf2* and *αB-crystallins genes*. This coupling manifests in the modulation of the lens and heart phenotypes of αB-crystallins loss-of-function zebrafish lines. Unexpectedly, Nrf2 deficiency, which increases the oxidative load, suppressed the lens defect phenotype in the *αBa-crystallin* knockout zebrafish embryos, but enhanced the heart edema phenotype characteristic of these lines. Transcriptome analysis identified distinct molecular pathways activated in response to impaired *nrf2* function in *αBa-crystallin* depleted lens and heart tissues. We found that Nrf2 deficiency activates cholesterol biosynthesis pathways in the *αBa-crystallin* mutated lens. In contrast, Nrf2 deficiency drives transcriptional change in the extracellular region and tight junction pathways in the heart tissue of *αBa-crystallin* mutant. To our knowledge, this is the first evidence of the role of sHSP in the oxidative stress response and sets the stage for an in-depth investigation of how transcriptional link to Nrf2 is mediated.

## Results

### Tissue-specific nrf2 regulation of αB-crystallin transcripts

To investigate how oxidative stress pathways modulate αB-crystallin expression and/or function, we utilized sensitized zebrafish lines bearing a mutation within *nrf2* that substantially reduces its activity [47]. The *nrf2^fh318^* zebrafish line has a mutation in the basic region of the DNA binding domain (exon 5), which leads to decreased transcriptional induction of Nrf2 target genes, including antioxidant enzymes. We observed that the mRNA level of *nrf2* downstream genes such as *gstp1* and *prdx1* were lower in the lens, and heart tissues of *nrf2*^fh318/fh318^ lines relative to WT fish (Fig S1). These results are consistent with previous studies although ours were carried out in the absence of oxidative stress [47]. We also confirmed that the level of *αBa-crystallin* and *αBb-crystallin* transcripts was relatively higher in the lens compared to heart and brain tissues, whereas the levels of *nrf2* transcripts in heart and brain was higher than in the lens (Fig S2).

To test if αB-crystallin expression is affected by Nrf2 deficiency, mRNA levels of *αBa-crystallin* and *αBb-crystallin* genes (hereafter, we use the nomenclature *“cryaba”* and *“cryabb”* for simplicity) were compared between WT and *nrf2*^fh318/fh318^ embryos by qRT-PCR analysis. Although the levels of *cryaba* transcript were not changed between WT and *nrf2*^fh318/fh318^, *cryabb* transcription was highly upregulated in Nrf2 deficient embryos (Fig 1A). This modulation of *cryabb* transcripts appears tissue specific. *cryaba* transcripts were not substantially changed in whole eyes, lens, heart, and brain tissues of *nrf2* mutated zebrafish relative to WT (Fig 1B). In contrast, *cryabb* transcripts were upregulated strongly in the heart and brain tissues of Nrf2 deficient zebrafish (Fig 1C, Fig S3). To test whether transcription of *cryabb* can be upregulated in response to increased oxidative stress, we measured the level *cryabb* mRNA after treating 4 days post-fertilization (dpf) WT embryos with 800 μM tert-Butyl hydroperoxide (tBHP) for 2 hours. We observed an increase in the *cryabb* transcript (~1.5 fold) with tBHP treatment (Fig S4B) although to a lesser extent compared to reduced activity of Nrf2.

**Figure 1.**
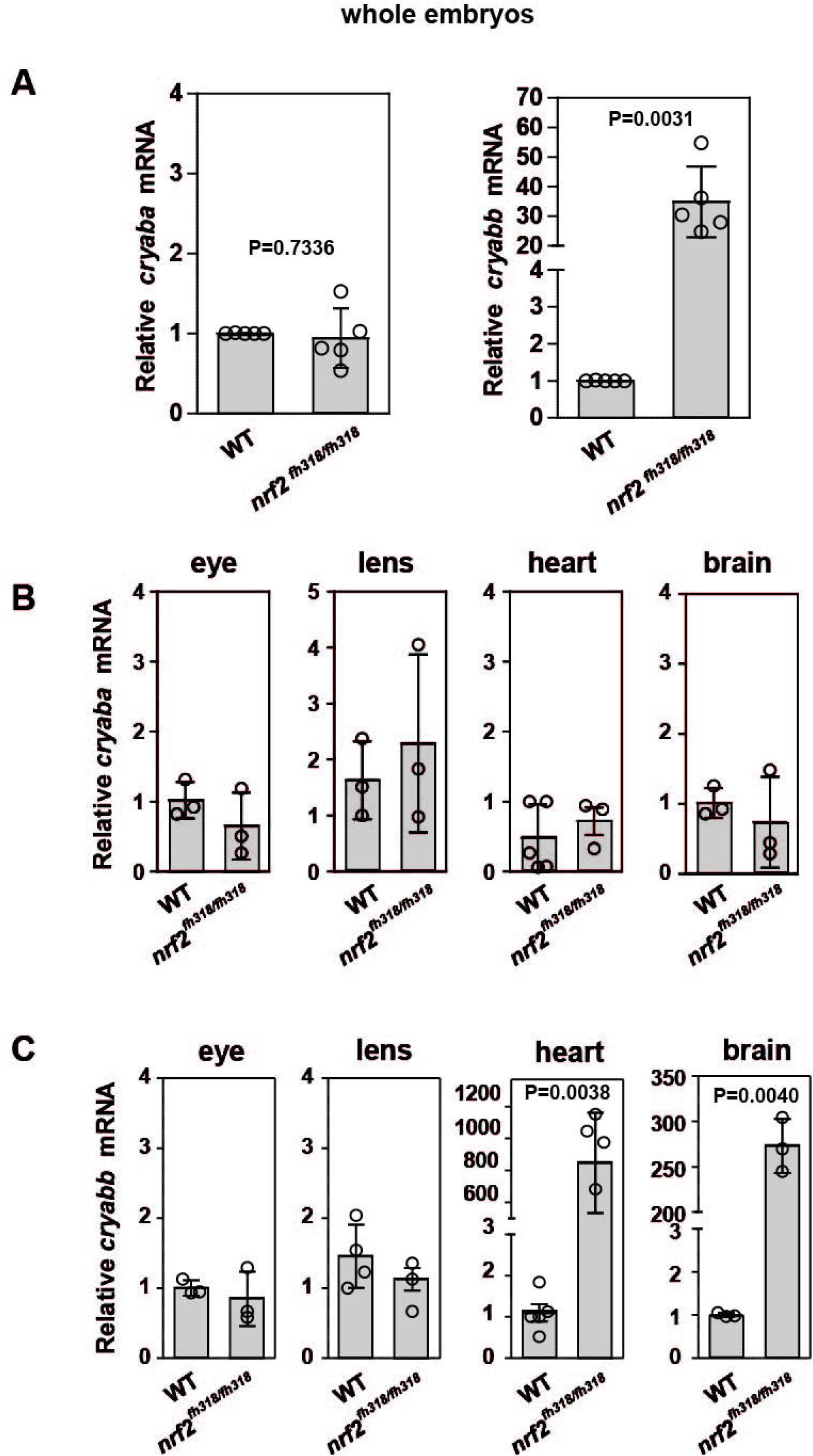
Zebrafish *cryab* transcripts are regulated in a tissue-dependent manner. **(A)** Relative expression of *cryaba* and *cryabb* genes was compared between WT and *nrf2*^fh318/fh318^ embryos at 4 dpf using qRT-PCR analysis. Data are expressed as mean ± SD from 5 independent measurements The relative mRNA levels of *cryaba* **(B)** and *cryabb* **(C)** were measured between WT and *nrf2*^fh318/fh318^ by qRT-PCR in the eyes, lens, heart, and brain tissues of 10-month-old zebrafish. Data are expressed as mean ± SD from at least *n*=3 for brain, eye, lens, and heart. *P*-values were calculated using two-tailed *t*-test

To investigate crosstalk between αB-crystallin and *nrf2*, we assessed the regulation of *nrf2* and *nrf2* downstream targets, including antioxidant genes, in αB-crystallin knockout lines (*cryaba^-/-^, cryabb^-/-^, cryaba^-/-^;cryabb^-/-^* double mutants). Although the level of *nrf2* mRNA was not significantly changed in WT and αB-crystallin knockout lines, its targets *gpx1a* and *prdx1* were upregulated in *cryabb^-/-^* embryos (Fig S5). Taken together, these results suggested a transcriptional link between *nrf2* and *cryabb* but not *cryaba*. Further detailed experiments are needed to examine the tissue-specific mechanisms underlying the upregulated antioxidant genes in the *cryabb^-/-^* lines.

### Nrf2 deficiency suppresses lens defects in the *cryaba^-/-^* mutant

In light of the transcriptional relationship described above, we explored its consequences on the phenotypic manifestation of the loss of αB-crystallin function. Previously, we showed that knockout of either αB-crystallin causes lens defects in zebrafish embryos [48, 49]. αB-crystallin mutant embryos, *cryaba^-/-^* and *cryabb^-/-^*, presented lens abnormalities at 4 dpf (Fig 2A). The phenotypic features characterizing these lines appeared as round puncta spread across the lens. These puncta lead to opacity and changes in light scattering. Consistent with our previous result, ~50% and ~30% of *cryaba^-/-^* and *cryabb^-/-^ embryos* exhibited lens abnormalities, respectively (Fig S6). In comparison, *nrf2^fh318/fh318^* showed slightly increased percentage of lens defects (~20%) relative to WT embryos (~10%) (Fig 2B).

**Figure 2.**
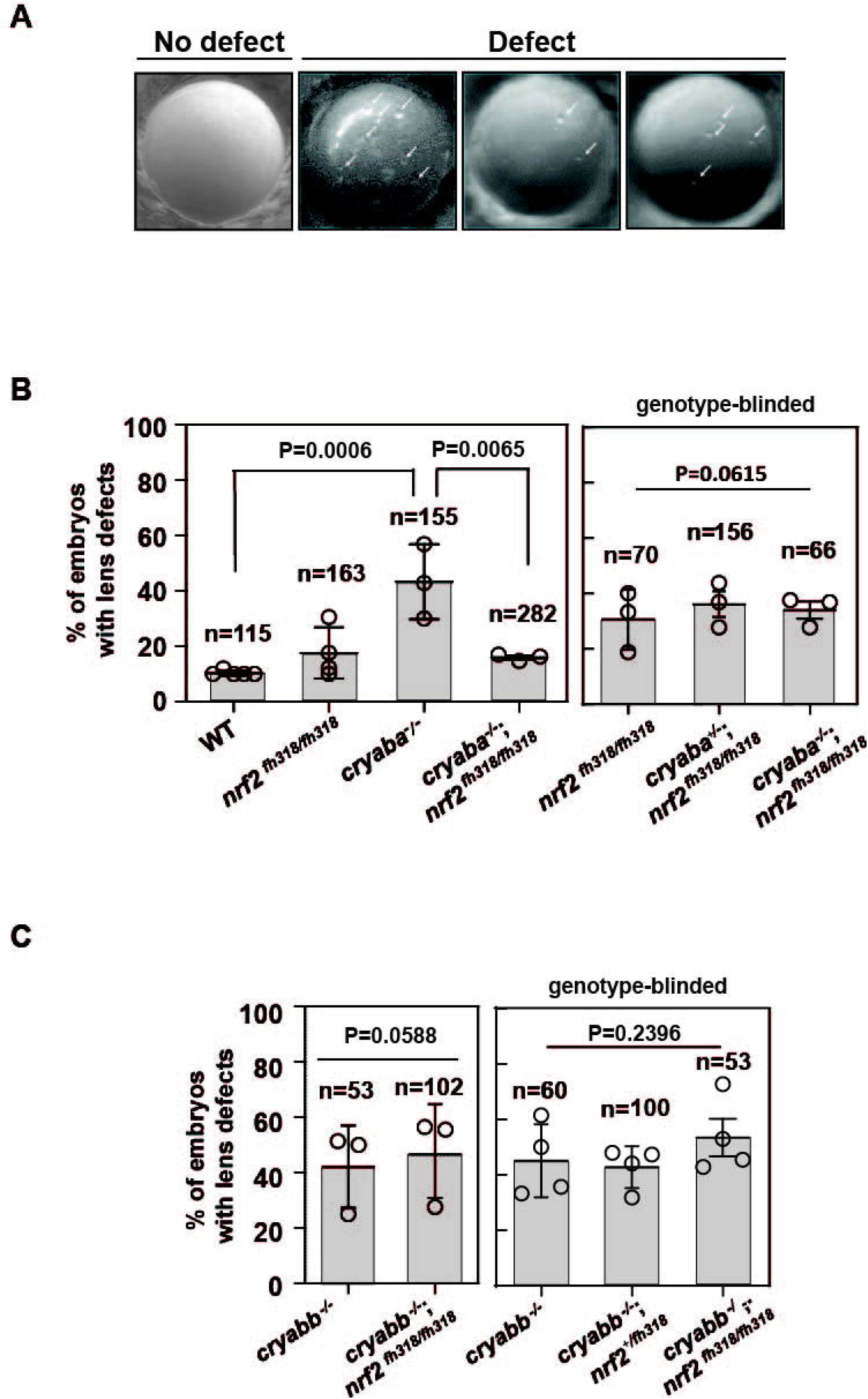
Nrf2 deficiency rescues lens defects in *cryaba^-/-^*, but not *cryabb*^-/-^. **(A)** Representative images of lens defects in zebrafish embryos at 4 day post fertilization (dpf). **(B, left panel)** The percentage of embryos showing lens defects for WT, *nrf2*^fh318/fh318^, *cryaba^-/-^* and *cryaba^-/-^*;*nrf2*^fh318/fh318^ was compared. **(B, right panel)** To confirm the previous results, embryos from *cryaba^+/-^*;*nrf2*^fh318/fh318^ incross were collected, and the lens defect of each embryo was measured in a genotype-blinded at 4 dpf. Then *cryaba* genotyping of individual embryos was determined to compare the percentage of lens defects between *nrf2*^fh318/fh318^ and *cryaba^-/-^*;*nrf2*^fh318/fh318^. **(C, left panel)** The percentage of lens defect between *cryaba^-/-^* and *cryaba^-/-^*;*nrf2*^fh318/fh318^ embryos. **(C, right panel)** Embryos from *cryaba^-/-^*;*nrf2*^fh318/+^ incross were collected, and lens abnormalities were examined before *nrf2* genotyping. Then the percentage of lens defects between *cryabb^-/-^* and *cryabb*^-/-^;*nrf2*^fh318/fh318^ was analyzed. Data are expressed as mean ± SD from at least three independent experiments. *P*-values were calculated using one-way ANOVA and two-tailed *t*-test.

To investigate how Nrf2 deficiency affects lens development in the absence of αB-crystallin, we generated αB-crystallin/Nrf2 double knockout lines (*cryaba*^+/-^;*nrf2*^fh318/fh318^, *cryaba*^-/-^;*nrf2*^fh318/fh318^, *cryabb*^-/-^;*nrf2*^fh318/+^, *cryabb*^-/-^;*nrf2*^fh318/fh318^). Then, we compared the penetrance of lens abnormalities in the progeny of WT, *nrf2*^fh318/fh318^, *cryaba^-/-^* and *cryaba*^-/-^;*nrf2*^fh318/fh318^ embryos. Strikingly, we found that the elevated level of lens defects in the *cryaba^-/-^* lines was suppressed in the *cryaba^-/-^*/*nrf2*^fh318/fh318^ (Fig 3B, left panel). To confirm this result, we in-crossed *cryaba*^+/-^;*nrf2*^fh318/fh318^ adult zebrafish and screened their embryos for lens defects in a genotype-blinded experimental protocol. After the lenses were imaged and sorted into normal and defective lens groups (Fig 2A), we categorized the *cryaba* genotype of individual embryos into three groups (*cryaba*^+/+^;*nrf2*^fh318/fh318^, *cryaba^+/-^*;*nrf2*^fh318/fh318^, *cryaba*^-/-^;*nrf2*^fh318/fh318^). The results (Fig 3B, right panel) are consistent with the conclusion that Nrf2 deficiency reduces the penetrance of the lens phenotype induced by αBa-crystallin loss of function. In contrast, we did not observe large changes in the percentage of lens defects between *cryabb^-/-^* and *cryabb*^-/-^;*nrf2*^fh318/fh318^ embryos, suggesting that the lens defects in *cryabb* KO lines cannot be rescued by abrogation of Nrf2 function (Fig 2C).

**Figure 3.**
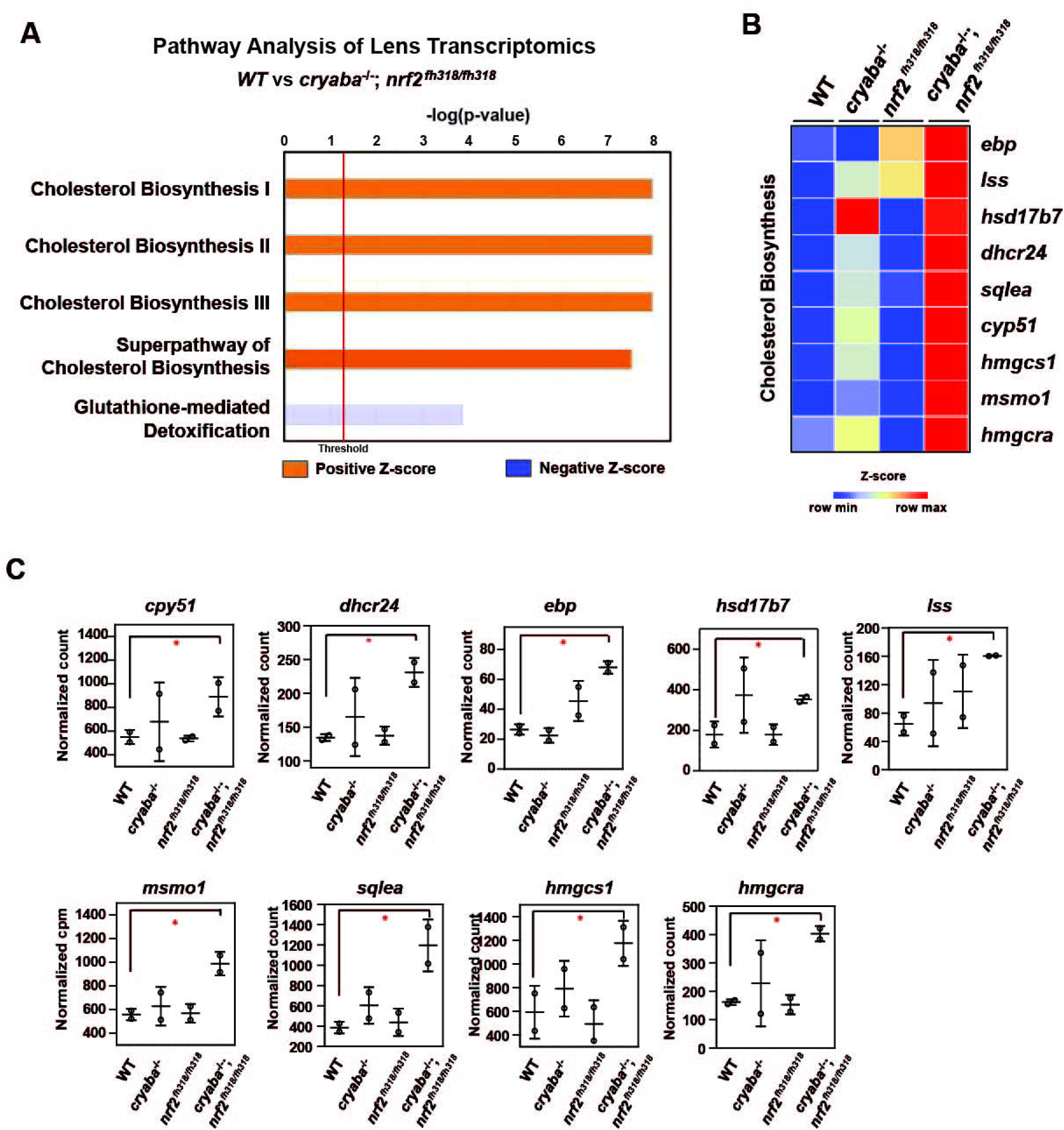
The cholesterol biosynthesis pathway is elevated in the lens of *cryaba^-/-^*;*nrf2*^fh318/fh318^. **(A)** Lens RNA-seq data were analyzed through ingenuity pathway analysis (www.ingenuity.com). Orange bars that cross the threshold line (*P* < 0.05) indicate upregulated pathways in the lens of *cryaba^-/-^*;*nrf2*^fh318/fh318^ compared to WT. **(B)** Heatmap of enriched genes in the superpathway of cholesterol biosynthesis. Z-scores were calculated for each gene, and these were plotted instead of the normalized expression values. **(C)** Bar charts represent the normalized count of each transcript in the cholesterol biosynthesis pathway in the lens tissues of WT, *nrf2*^fh318/fh318^, *cryaba^-/-,^* and *cryaba^-/-^*;*nrf2*^fh318/fh318^. * Indicate False Discovery Rate (FDR) < 0.05.

### Upregulation of cholesterol biosynthesis pathway in the lens of cryaba^-/-^;nrf2^fh318/fh318^ double mutant lines

To identify the molecular pathway(s) mediating the suppression of lens defects in the *cryaba* mutant by Nrf2 deficiency, we performed high-throughput transcriptome profiling on adult zebrafish lens tissues from the WT, *cryaba^-/-^, nrf2*^fh318/fh318^, and *cryaba*^-/-^;*nrf2*^fh318/fh318^ lines. Differentially expressed (DE) genes between groups were determined using Illumina’s Dragen pipeline. Then, the DE genes were processed using Ingenuity Pathway Analysis (IPA) to define the significantly regulated signaling pathways [50]. Interestingly, cholesterol biosynthesis was identified as the primary upregulated pathway in the lens of *cryaba^-/-^*;*nrf2*^fh318/fh318^ compared to WT (Fig 3A). mRNA of genes encoding enzymes involved in the superpathway of cholesterol biosynthesis, *cyp51, dhcr24, ebp, lss, msmo1, sqlea, hmgcra, and hmgcs1*, were upregulated in the lens of the *cryaba*^-/-^;*nrf2*^fh318/fh318^ line compared to WT (Fig 3B, C).

Based on the RNA-seq data, we hypothesized a correlation between activated cholesterol biosynthesis and the alleviated lens defects in the *cryaba*^-/-^;*nrf2*^fh318/fh318^ line. To test this hypothesis, we investigated whether the lens of *cryaba*^-/-^;*nrf2*^fh318/fh318^ embryos exhibits lower tolerance to treatment with statins, which are competitive inhibitors of the HMG-CoA-R enzyme, an early rate-limiting step in cholesterol synthesis [51]. For this purpose, the *cryaba*^-/-^;*nrf2*^fh318/fh318^ embryos were challenged with two different statins, atorvastatin, and lovastatin, to inhibit the sterol biosynthetic pathway. *cryaba^-/-^*;*nrf2*^fh318/fh318^ embryos, incubated with 5 μM atorvastatin from 1 to 4 dpf, showed increased lens abnormalities (Fig 4A). In contrast, vehicle-treated *cryaba*^-/-^;*nrf2*^fh318/fh318^ (DMSO control) or WT embryos incubated with the same regimen of atorvastatin remained predominantly normal. In addition, we noticed that atorvastatin-exposed *cryaba*^-/-^;*nrf2*^fh318/fh318^ embryos have significantly reduced lens area than vehicle-treated control (Fig 4B). Similarly, *cryaba*^-/-^;*nrf2*^fh318/fh318^ embryos displayed a higher percentage of lens defects than WT in response to 4 μM lovastatin (Fig 4C). Together, these results suggest that upregulation of the cholesterol synthesis pathway in the lens is a plausible mechanism involved in the suppression of lens abnormalities in the *cryaba*^-/-^;*nrf2*^fh318/fh318^.

**Figure 4.**
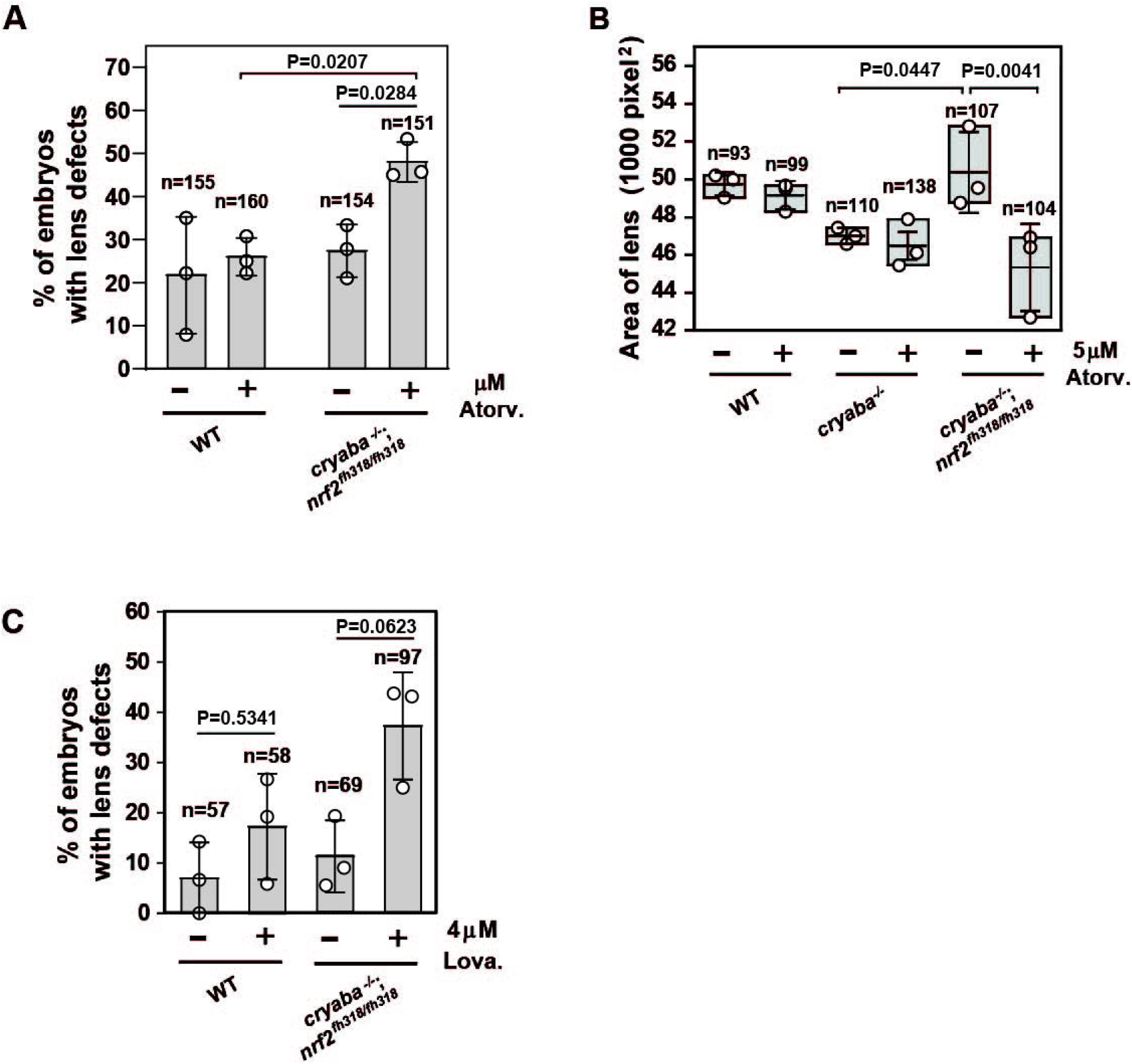
Increased penetrance of lens defects in *cryaba^-/-^*;*nrf2*^fh318/fh318^ in response to treatment of statins. **(A)** Zebrafish embryos were treated with either vehicle (1% DMSO) or 5 μM atorvastatin from 36 hours post-fertilization (hpf) until 4 dpf. Percentage of embryos showing lens defects between WT and *cryaba^-/-^*;*nrf2*^fh318/fh318^ were compared at 4 dpf. **(B)** The size of the lens of WT, *cryaba^-/-^*, and *cryaba^-/-^*;*nrf2*^fh318/fh318^ embryos was measured in the presence and absence of 5 μM atorvastatin treatment. **(C)** WT and *cryaba^-/-^*;*nrf2*^fh318/fh318^ embryos were treated with 4 μM lovastatin for 16 hours before examining lens abnormalities at 4 dpf. Data are expressed as mean ± SD from three independent experiments. Statistical significance was calculated using two-way ANOVA.

### *Dexamethasone-induced cardiac edema of* Nrf2 deficient *Zebrafish is potentiated by αB-crystallin deficiency*

In addition to lens defects, loss of αB-crystallin function is associated with a cardiac phenotype that presents as embryonic cardiac edema [48]. The penetrance of this phenotype increases in response to stress induced by exposure to external glucocorticoid receptor agonists such as dexamethasone (Dex) as we have previously described [48]. Because it has been reported that Nrf2 has a protective role in cardiac cells under oxidative stress [52], we examined whether the stress-induced heart edema of αB-crystallin KO lines is modulated by Nrf2 deficiency. For this purpose, we compared the heart areas (see methods section) of zebrafish embryos derived from the loss-of-function lines of αB-crystallins, Nrf2, and crosses thereof *cryaba^-/-^*;*nrf2*^fh318/fh318^ and *cryabb^-/-^*;*nrf2*^fh318/fh318^. All single KO lines and *nrf2*^fh318/fh318^, treated with 50 μM dexamethasone (Dex) from 1 to 4 dpf, developed pericardial edema manifested by increased heart area relative to WT embryos (Fig 5A-C). Remarkably, we observed a large increase in the penetrance of the phenotype in the *cryaba*^-/-^;*nrf2*^fh318/fh318^ line that was further accentuated in the presence of Dex. (Fig 5B, D). Relative to *cryaba*^-/-^;*nrf2*^fh318/fh318^, cardiac edema was blunted in *cryabb*^-/-^;*nrf2*^fh318/fh318^ embryos treated with Dex. Furthermore, the vehicle-treated *cryabb*^-/-^;*nrf2*^fh318/fh318^ group retained a WT-like distribution (Fig 5C).

**Figure 5.**
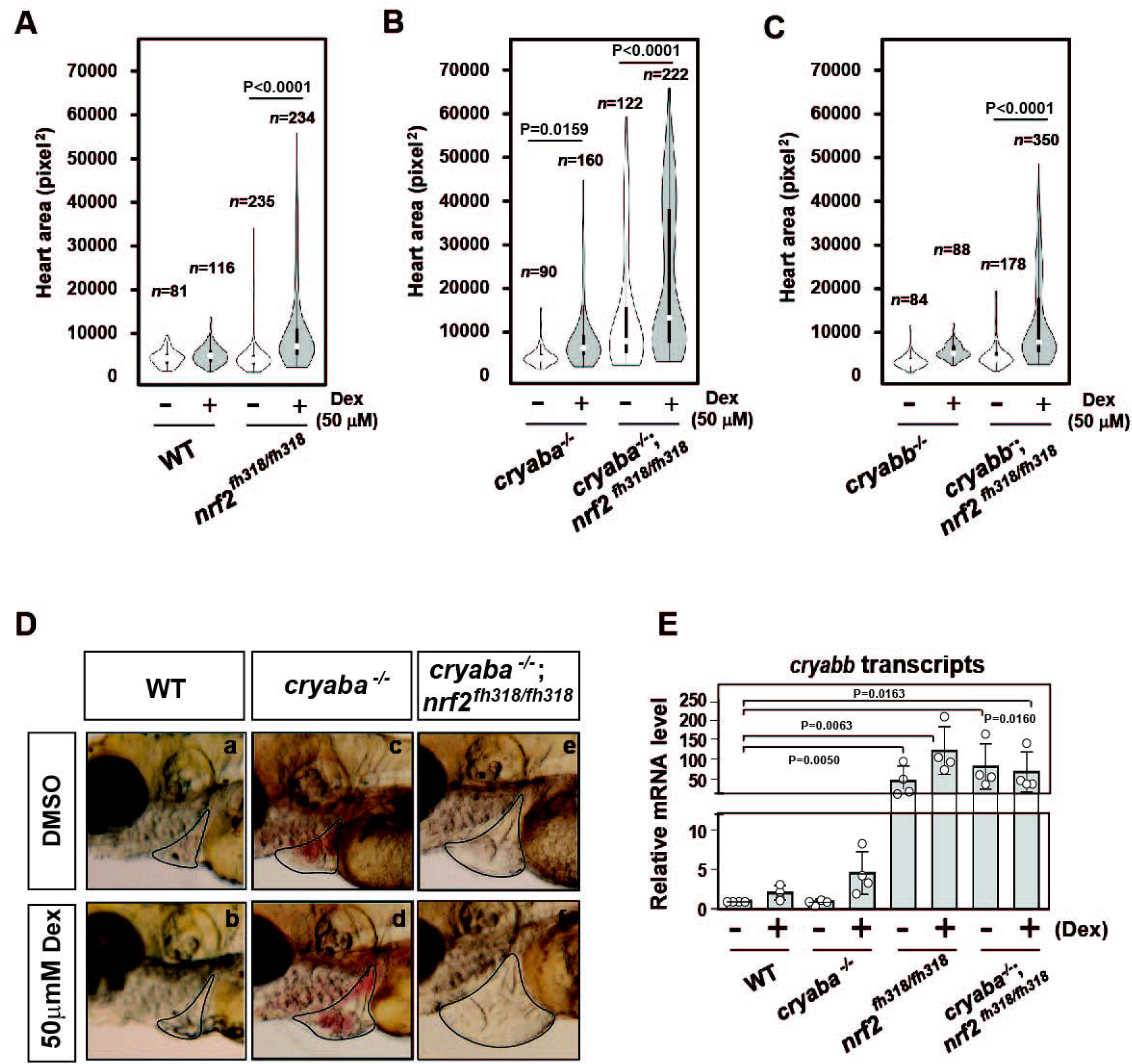
Nrf2 deficiency aggravates the heart edema phenotype of *cryaba^-/-^* fish in response to dexamethasone (Dex). The heart area of *nrf2*^fh318/fh318^ embryos was measured following treatment with 50 μM Dex and compared to **(A)** the WT embryos, **(B)** *cryaba^-/-^* and *cryaba^-/-^*;*nrf2*^fh318/fh318^, and **(C)** *cryaba^-/-^* and *cryaba^-/-^*;*nrf2*^fh318/fh318^ following the same treatment regimen. **(D)** Representative images of cardiac phenotypes in WT, *cryaba^-/-^* and *cryaba^-/-^*;*nrf2*^fh318/fh318^ treated with either vehicle (DMSO) or Dex. **(E)** Relative changes in *cryabb* mRNA in 4 dpf embryos as measured by qRT-PCR. Data are expressed as mean ± SD obtained from three independent experiments. Statistical significance was calculated using two-way ANOVA.

Because the discovery of *cryabb* mRNA upregulation in Nrf2 deficient heart tissues (Fig 1C) suggests *cryabb* is transcriptionally activated in response to oxidative stress, we further explored the level of *cryabb* transcript in the presence of Dex. The transcription of *cryabb* mRNA appears to be upregulated with Dex treatment in WT (~2-fold) and *cryaba* KO (~3-fold) at 4 dpf (Fig 5E) although the p values for two-way ANOVA were higher than 0.05. Furthermore, *nrf2*^fh318/fh318^ and *cryaba^-/-^*;*nrf2*^fh318/fh318^ embryos showed strongly upregulated *cryabb* mRNA regardless of the presence or absence of Dex (Fig 5E). Taken together, these results suggest that the two zebrafish αB-crystallin orthologues, *cryaba* and *cryabb*, have different regulatory mechanisms in the heart in response to oxidative stress. Our data suggest that *cryabb* functions as a stress-response gene regulated by both glucocorticoid stress and oxidative stress. Further detailed experiments are needed to pinpoint the mechanism of the transcriptional control *cryabb* under different forms of stress.

### Changes in the extracellular region and tight junction pathways in the heart of cryaba^-/-^;nrf2^fh318/fh318^

To gain mechanistic insight into the origin of the heart edema phenotype, we performed transcriptome analysis of heart tissues from adult zebrafish comparing WT, *cryaba*^-/-^, *nrf2*^fh318/fh318^, and cryaba^-/-^;*nrf2*^fh318/fh318^. RNA-seq analysis identified DE genes in the heart tissue of Nrf2 deficient zebrafish (*nrf2*^fh318/fh318^ and *cryaba*^-/-^;*nrf2*^fh318/fh318^), including upregulation of *cryabb* transcripts, which is consistent with our data using qRT-PCR analysis (Fig 1C, Fig S7F). To derive a global understanding of how these genes affect heart health, the DE genes with a false discovery rate (FDR) cut off ≤0.05 between groups were used for further pathway analysis. The most significant gene ontology (GO) was calculated by using the WEB-based GEne SeT AnaLysis (WebGestalt) Toolkit [53]. The top important GO terms between *cryaba*^-/-^;*nrf2*^fh318/fh318^ *versus* WT included the extracellular region, supermolecular fiber, and bicellular tight junction (Fig6A). The extracellular region and tight junction pathways were also listed in the most enriched GO cluster between *cryaba*^-/-^;*nrf2*^fh318/fh318^ *versus cryaba*^-/-^ (Fig6B). These findings suggest a significant alteration of genes related to extracellular interactions in the heart of *cryaba^-/-^*;*nrf2*^fh318/fh318^ embryos.

**Figure 6.**
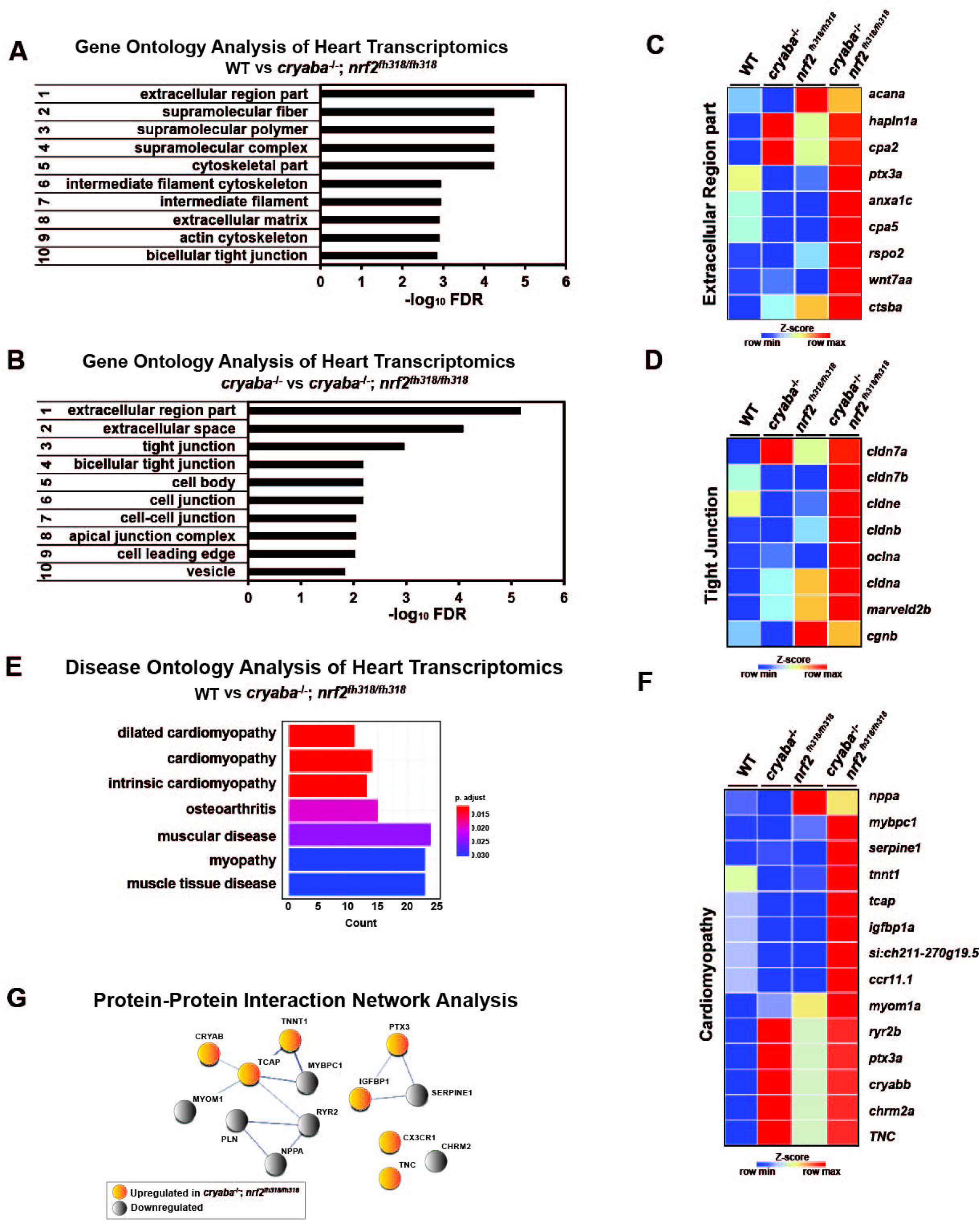
Transcriptome analysis reveals changes in genes belonging to the extracellular region in the heart of *cryaba^-/-^*;*nrf2*^fh318/fh318^. The top-ranked significant GO clusters in heart tissues between **(A)** WT versus *cryaba^-/-^*;*nrf2*^fh318/fh318^, and **(B)** *cryaba^-/-^* versus *cryaba^-/-^*;*nrf2*^fh318/fh318^. Heatmaps illustrate the transcripts in **(C)** the extracellular region GO term and **(D)** tight junction GO term. **(E)** The top terms in the Disease Ontology (DO) enrichment analysis highlight the number of genes enriched in each term (x-axis). The adjusted *P*-value of each term is indicated by color according to the legend. **(F)** The enriched genes in the cardiomyopathy DO cluster are illustrated by a heatmap. **(G)** Interaction between human homologs of transcripts in the cardiomyopathy DO term was assessed using STRING tool [55].

To identify the extracellular components which are exclusively changed in the heart tissue of *cryaba^-/-^*;*nrf2*^fh318/fh318^, the subset of genes in the GO term within the extracellular region from each comparison (*cryaba*^-/-^;*nrf2*^fh318/fh318^ *vs*. WT, and *cryaba^-/-^*;*nrf2*^fh318/fh318^ *vs*. *cryaba*^-/-^) was sorted by Venn-diagram analysis (Fig S8, Table S3, 4). The result identified *cpa5, cstba, cpa2, anxa1c, hapln1a, ptx3a, rspo2, wnt7aa*, and *acana* genes which are closely involved in the degradation, remodeling, or inflammation of extracellular matrix (ECM) (Fig 6C). In addition, the transcripts which encode components in the tight junction such as *cldn7b, cldn3, cldnb, oclna, cagnb*, and *marveld2b*, showed higher expression in the heart tissues of *cryaba*^-/-^;*nrf2*^fh318/fh318^ than *cryaba^-/-^* (Fig 6D, Fig S9). These results suggest that molecular changes of ECM and tight junction pathways in the heart in *cryaba^-/-^*;*nrf2*^fh318/fh318^ correlate with greater penetrance of the cardiac edema phenotype.

Finally, we performed Disease Ontology (DO) analysis with known disease markers to explore the potential pathological mechanism underlying the enhanced heart edema in *cryaba^-/-^*;*nrf2*^fh318/fh318^. For this purpose, the DE genes between *cryaba^-/-^*;*nrf2*^fh318/fh318^ and WT in the heart tissues were analyzed using DOSE [54] to calculate enrichment of DO terms. The results confirmed that the transcriptional changes in the heart of *cryaba^-/-^*;*nrf2*^fh318/fh318^ are significantly associated with muscle diseases, specifically cardiomyopathy (Fig 6E, F). We also noticed that the human homologs of transcripts in the cardiomyopathy DO are functionally clustered via STRING, a database of protein-protein interactions [55] (Fig 6G). This result further supports the transcriptional changes in *cryaba^-/-^*;*nrf2*^fh318/fh318^ are functionally associated with heart health.

## Discussion

### zebrafish *αB-crystallin* genes are regulated under oxidative stress in a tissue-specific manner

Despite their ubiquitous expression, the physiological roles of sHSP, beyond chaperone activity, continue to be enigmatic. While they have been implicated in numerous stress responses, there has been a lack of systematic investigation of how they are coupled to cellular protective pathways. To address this outstanding question, we have utilized zebrafish as a tractable model system to dissect the role of αB-crystallin both in the lens and other tissues. In previous work, we demonstrated that αB-crystallins are critical for lens development and that the loss of function of αB-crystallins compromises the resistance of the heart to stress [48, 49]. Here, we explored the role of the two αB-crystallins in response to oxidative stress signaling. The finding of tissue-specific transcriptional link between *nrf2* and *cryabb* but not *cryaba* in zebrafish is novel and is consistent with the recent findings that Nrf2 can function as a repressor [56, 57]. Equally novel is the identification of molecular pathways modulated in loss-of-function zebrafish lines. The results presented here suggest that *cryabb* plays a role in the stress response in brain and heart tissues, while *cryaba* mainly has a function limited to the lens in zebrafish, most likely as a chaperone. Overall, our results set the stage for further studies of how *nrf2* and *cryabb* are coupled in the zebrafish lens and heart. In both tissues, long-lived cells experience sustained oxidative loads that lead to a number of pathologies.

### A novel intersection between lens integrity and cholesterol biosynthesis pathway

A major finding of this paper is that lens defects in the *cryaba^-/-^* line were rescued in *cryaba^-/-^*;*nrf2*^fh318/fh318^ line(Fig 2). Furthermore, RNA-seq analysis correlated this rescue with the upregulation of the cholesterol biosynthesis pathway (Fig 3). Following up on the hypothesis of a connection between cholesterol biosynthesis and alleviated lens defects in *cryaba^-/-^*;*nrf2*^fh318/fh318^, we utilized statins, HMG-CoA reductase inhibitors, to lower cholesterol synthesis. Indeed, we determined that treatment with atorvastatin or lovastatin increased the penetrance of lens defect in *cryaba^-/-^*;*nrf2*^fh318/fh318^.

The beneficial role of high cholesterol in the lens has been described and specifically linked to a reduction in oxygen transport across membranes to protect against oxidative damage [58–60]. Moreover, elevating cholesterol precursors, particularly lanosterol, have been studied as a means to reverse protein aggregation in cataracts [58, 59, 61, 62]. In this context, a genetic mutation in lanosterol synthase (*lss*) is associated with cholesterol deficiency-associated cataracts [63]. However, the precise molecular mechanism linking cholesterol biosynthesis and cataract remains largely unknown. Here, we provide an additional connection between cholesterol and lens integrity using high-throughput transcriptome profiling. Given that lens RNA-seq analysis revealed upregulation of cholesterol biosynthetic enzymes, we propose that lathosterol is likely the upregulated sterol in the lens, although the means by which Nrf2 deficiency upregulates cholesterol biosynthetic enzymes is still unclear. Thus, further experiments are needed to delineate the intersection between the cholesterol synthesis pathway and Nrf2 in the lens.

Our results are consistent with previous reports have uncovered Nrf2-dependent regulation of cholesterol biosynthesis in mouse liver and in cultured liver cells. A pharmacogenomics investigation of the Nrf2 activator 3H-1,2-dithiole-3-thione (D3T) in mice showed RNA enrichment of enzymes involved in cholesterol biosynthesis whereas no effect on these enzymes was observed as a consequence of treatment with the Nrf2 activator 1-[2-cyano-3,12-dioxooleana-1,9(11)-dien-28-oyl]-imidazole (CDDO-Im) [64]. In contrast, a dietary supplement found to activate Nrf2 downregulated the cholesterol biosynthesis pathway [65]. The contrasting results notwithstanding, these studies confirm a relationship between Nrf2 and cholesterol biosynthesis pathway and point out to the need for a more thorough understanding of how Nrf2 regulates this pathway.

### Loss of nrf2 function increases the penetrance of the cardiac phenotype in cryaba^-/-^ but not cryabb^-/-^ zebrafish

In contrast to the lens phenotype, the fraction of embryos displaying stress-induced heart edema in *cryaba^-/-^* was increased by the loss of Nrf2 function. More importantly, a higher penetrance of the heart defect phenotype was observed in *cryaba^-/-^*;*nrf2*^fh318/fh318^ embryos even without Dex treatment (Fig 5B). Furthermore, the search for disease-associated molecular pathways revealed cardiomyopathy as the top DO cluster in *cryaba^-/-^*;*nrf2*^fh318/fh318^ adult zebrafish heart, suggesting a detrimental synergetic effect of the depletion of *nrf2* and *cryaba* in heart tissue.

While more studies are needed to mechanistically understand the link between these various pathways, our findings can be framed in the extensive literature that implicates αB-crystallin in the oxidative balance of cardiomyocytes in mouse lines as well as in humans [66–68]. First identified in investigations of the human cardiomyopathy mutant R120G of αB-crystallin, subsequent studies have elaborated a link between αB-crystallin and reductive stress. Zebrafish is an ideal model to investigate this link further.

## Experimental Procedures

### Zebrafish maintenance and breeding

AB wild-type strain zebrafish (Danio rerio) were used. The embryos were obtained by natural spawning and raised at 28.5 °C on a 14:10 h light/dark cycle in egg water 30 mg/L instant ocean in deionized water containing 0.003% PTU (w/v) to prevent pigment formation. Embryos were staged according to their ages (in dpf). The following mutant and transgenic fish lines were used: *cryaba*^vu612^ (*cryαba^-/-^*); *cryabb*^vu613^ (*cryαbb^-/-^*); *nrf2*^fh318^. All animal procedures were approved by the Vanderbilt University Institutional Animal Care and Use Committee.

### Quantitative reverse transcription PCR

Zebrafish were sacrificed, and lens, heart, and brain tissues were dissected as described [69]. Tissues were immediately snap-frozen in liquid nitrogen, and RNA was extracted from all samples simultaneously using TRIzol (Invitrogen) and RNA clean & concentrator kit (Zymo Research). 500 μg of total RNA was then used as a template with the SuperScript III First-Strand Synthesis kit (Invitrogen) to produce cDNA. The specific targets were amplified by RT-PCR using oligonucleotides in supplementary tables 1 and 2. *β-actin* is used as the internal control. ANOVA and t-tests were performed to calculate the *P*-value to determine the significant difference between samples.

### RNA-Seq

Total RNA from zebrafish tissues was isolated simultaneously using TRIzol (Invitrogen) and RNA clean & concentrator kit (Zymo Research). RNA-Seq libraries (n=2) were processed at the Vanderbilt Technologies for Advanced Genomics (VANTAGE) core. Briefly, samples were processed using the TruSeq Standard sample Prep Kit (Illumina) to prepare cDNA libraries after Poly(A) selection. The libraries were sequenced on an Illumina NovaSeq 6000 to a depth of 50 million at 150 bp paired-end reads per library. For lens RNA-seq, FASTQ reads were aligned through Illumina’s Dragen RNA Seq pipeline. EdgeR (3.30.3) packages were used to measure differential gene expression with genes that achieved a count per million mapped reads (CPM). For heart RNA-seq, Advanced Genomics Analysis and Research Design (VANGARD) perform DEG and further analysis.

### Gene and Disease ontology

For lens RNA seq, DEG (FDR cutoff ≤ 0.01) was analyzed through the use of IPA (QIAGEN, https://www.qiagenbioinformatics.com/products/ingenuitypathway-analysis). The GO clusters with significant *P*-value were taken for further analysis. To use the human Disease Ontology database, the zebrafish genes were first matched to their human orthologues using Ensembl biomart (www.ensembl.org/biomart). Then, the gene set of human orthologues was processed using Bioconductor DOSE packages [54] with a *P*-value ≤0.05 cutoff to identify corresponding DO terms.

### Microscopy and image processing

Lenses of live embryos in 0.3X Danieau water with PTU/tricaine were analyzed by bright field microscopy (Zeiss Axiozoom V16) at 4 dpf. The percent of lens defects was scored as defined in our previous study [48]. Briefly, the phenotypic characteristic manifested as a spherical, shiny droplet spreading across the lens, which was classified as a defect. For heart imaging, the heart area of live embryos at 4 dpf was analyzed by bright field Axiozoom microscopy. Quantification of the size of the heart area was performed by ImageJ [70, 71].

### Drug treatments

1 dpf embryos were manually dechorionated, and 10 embryos were placed in one well of a 24-well plate (polystyrene, tissue culture grade) with 1 ml of 0.3x Danieau water. 50 μM dexamethasone (Sigma, D1756) diluted in 0.3x Danieau water was treated from 1 until 4 dpf to examine the cardiac phenotypes. For stain experiments, 2.5 or 5μM atorvastatin (Santa Cruz Biotechnology, sc-337542A) was treated from 1 to 4-dpf to observe lens defects, and 4 μM lovastatin (Santa Cruz) Biotechnology, sc-200850A) was treated for 16 hours before examining lens abnormalities at 4 dpf.

### Statistics

Statistical analyses were carried out with GraphPad Prism software 9 (Graphpad) by means of Student *t*-test or ANOVA. Comparison between groups were performed with Bonferroni or Tukey test for one-way or two-way ANOVA. Statistical significance was defined as *P* < 0.05

## Data Availability

RNA-seq data have been deposited in the ArrayExpress database at EMBL-EBI (www.ebi.ac.uk/arrayexpress) under accession number E-MTAB-12172.

## Supporting information

This article contains supporting information.

## Conflict of interest

The authors declare that they have no conflicts of interest with the contents of this article.

## Acknowledgements

The authors wish to thank Dr. Derek P. Claxton for a critical reading of the manuscript and helpful discussions.

## FOOTNOTES

This work was supported by EY12018 to HSM from the National Institutes of Health.

## The abbreviations used are

sHSP: small heat shock protein
dpf: days post-fertilization
qRT: quantitative Reverse Transcription
PTU: 1-phenyl-2-thiourea
Dex: dexamethasone
tBHP: tert-Butyl hydroperoxide
GO: Gene ontology
RNA-seq: RNA-sequencing
Atorv: atorvastatin
Lova: Lovastatin
FDR: False Discovery Rate

## Notes

### Competing Interest Statement

The authors have declared no competing interest.

## References

1. Truscott, R.J., Age-related nuclear cataract-oxidation is the key. Exp Eye Res, 2005. 80(5): p. 709–25.

2. Bartz, R.R., H.B. Suliman, and C.A. Piantadosi, Redox mechanisms of cardiomyocyte mitochondrial protection. Front Physiol, 2015. 6: p. 291.

3. Auten, R.L. and J.M. Davis, Oxygen toxicity and reactive oxygen species: the devil is in the details. Pediatr Res, 2009. 66(2): p. 121–7.

4. Tonelli, C., I.I.C. Chio, and D.A. Tuveson, Transcriptional Regulation by Nrf2. Antioxid Redox Signal, 2018. 29(17): p. 1727–1745.

5. Yamamoto, M., T.W. Kensler, and H. Motohashi, The KEAP1-NRF2 System: a Thiol-Based Sensor-Effector Apparatus for Maintaining Redox Homeostasis. Physiol Rev, 2018. 98(3): p. 1169–1203.

6. Vetter, C.J., et al., Cumulative deamidations of the major lens protein gammaS-crystallin increase its aggregation during unfolding and oxidation. Protein Sci, 2020. 29(9): p. 1945–1963.

7. Luo, C., et al., New insights into change of lens proteins’ stability with ageing under physiological conditions. Br J Ophthalmol, 2021.

8. Kopylova, L.V., et al., Age-related changes in the water-soluble lens protein composition of Wistar and accelerated-senescence OXYS rats. Mol Vis, 2011. 17: p. 1457–67.

9. Moreau, K.L. and J.A. King, Protein misfolding and aggregation in cataract disease and prospects for prevention. Trends Mol Med, 2012. 18(5): p. 273–82.

10. Duerksen, R., et al., Cataract blindness in Paraguay--results of a national survey. Ophthalmic Epidemiol, 2003. 10(5): p. 349–57.

11. Dunzhu, S., et al., Blindness and eye diseases in Tibet: findings from a randomised, population based survey. Br J Ophthalmol, 2003. 87(12): p. 1443–8.

12. Murthy, G.V., et al., A population-based eye survey of older adults in a rural district of Rajasthan: I. Central vision impairment, blindness, and cataract surgery. Ophthalmology, 2001. 108(4): p. 679–85.

13. Thulasiraj, R.D., et al., Blindness and vision impairment in a rural south Indian population: the Aravind Comprehensive Eye Survey. Ophthalmology, 2003. 110(8): p. 1491–8.

14. Zhao, J., et al., Prevalence of vision impairment in older adults in rural China: the China Nine-Province Survey. Ophthalmology, 2010. 117(3): p. 409–16, 416 e1.

15. Zheng, Y., et al., Prevalence and causes of visual impairment and blindness in an urban Indian population: the Singapore Indian Eye Study. Ophthalmology, 2011. 118(9): p. 1798–804.

16. Beebe, D.C., N.M. Holekamp, and Y.B. Shui, Oxidative damage and the prevention of age-related cataracts. Ophthalmic Res, 2010. 44(3): p. 155–65.

17. Shui, Y.B., et al., Oxygen distribution in the rabbit eye and oxygen consumption by the lens. Invest Ophthalmol Vis Sci, 2006. 47(4): p. 1571–80.

18. Wu, S.Y. and M.C. Leske, Antioxidants and cataract formation: a summary review. Int Ophthalmol Clin, 2000. 40(4): p. 71–81.

19. Garner, M.H. and A. Spector, Selective oxidation of cysteine and methionine in normal and senile cataractous lenses. Proc Natl Acad Sci U S A, 1980. 77(3): p. 1274–7.

20. Garner, M.H.and A. Spector, Sulfur oxidation in selected human cortical cataracts and nuclear cataracts. Exp Eye Res, 1980. 31(3): p. 361–9.

21. Truscott, R.J. and R.C. Augusteyn, The state of sulphydryl groups in normal and cataractous human lenses. Exp Eye Res, 1977. 25(2): p. 139–48.

22. Truscott, R.J.W. and M.G. Friedrich, Molecular Processes Implicated in Human Age-Related Nuclear Cataract. Invest Ophthalmol Vis Sci, 2019. 60(15): p. 5007–5021.

23. Giblin, F.J., Glutathione: a vital lens antioxidant. J Ocul Pharmacol Ther, 2000. 16(2): p. 121–35.

24. Liu, X.F., et al., Nrf2 as a target for prevention of age-related and diabetic cataracts by against oxidative stress. Aging Cell, 2017. 16(5): p. 934–942.

25. Rowan, S., et al., Aged Nrf2-Null Mice Develop All Major Types of Age-Related Cataracts. Invest Ophthalmol Vis Sci, 2021. 62(15): p. 10.

26. von Otter, M., et al., Nrf2-encoding NFE2L2 haplotypes influence disease progression but not risk in Alzheimer’s disease and age-related cataract. Mech Ageing Dev, 2010. 131(2): p. 105–10.

27. Bassnett, S., On the mechanism of organelle degradation in the vertebrate lens. Exp Eye Res, 2009. 88(2): p. 133–9.

28. Horwitz, J., Alpha-crystallin can function as a molecular chaperone. Proc Natl Acad Sci U S A, 1992. 89(21): p. 10449–53.

29. Kumar, P.A. and G.B. Reddy, Modulation of alpha-crystallin chaperone activity: a target to prevent or delay cataract? IUBMB Life, 2009. 61(5): p. 485–95.

30. Kaiser, C.J.O., et al., The structure and oxidation of the eye lens chaperone alphaA-crystallin. Nat Struct Mol Biol, 2019. 26(12): p. 1141–1150.

31. Slingsby, C., G.J. Wistow, and A.R. Clark, Evolution of crystallins for a role in the vertebrate eye lens. Protein Sci, 2013. 22(4): p. 367–80.

32. Sun, Z., et al., Mutations in crystallin genes result in congenital cataract associated with other ocular abnormalities. Mol Vis, 2017. 23: p. 977–986.

33. Berry, V., et al., The genetic landscape of crystallins in congenital cataract. Orphanet J Rare Dis, 2020. 15(1): p. 333.

34. Bhat, S.P. and C.N. Nagineni, alpha B subunit of lens-specific protein alpha-crystallin is present in other ocular and non-ocular tissues. Biochem Biophys Res Commun, 1989. 158(1): p. 319–25.

35. Klemenz, R., et al., Alpha B-crystallin is a small heat shock protein. Proc Natl Acad Sci U S A, 1991. 88(9): p. 3652–6.

36. Swist, S., et al., Maintenance of sarcomeric integrity in adult muscle cells crucially depends on Z-disc anchored titin. Nat Commun, 2020. 11(1): p. 4479.

37. Franssen, C., et al., alpha-B Crystallin Reverses High Diastolic Stiffness of Failing Human Cardiomyocytes. Circ Heart Fail, 2017. 10(3): p. e003626.

38. Chis, R., et al., alpha-Crystallin B prevents apoptosis after H2O2 exposure in mouse neonatal cardiomyocytes. Am J Physiol Heart Circ Physiol, 2012. 303(8): p. H967–78.

39. Bova, M.P., et al., Mutation R120G in alphaB-crystallin, which is linked to a desmin-related myopathy, results in an irregular structure and defective chaperone-like function. Proc Natl Acad Sci U S A, 1999. 96(11): p. 6137–42.

40. Boelens, W.C., Cell biological roles of alphaB-crystallin. Prog Biophys Mol Biol, 2014. 115(1): p. 3–10.

41. Kannan, S., et al., Nrf2 deficiency prevents reductive stress-induced hypertrophic cardiomyopathy. Cardiovasc Res, 2013. 100(1): p. 63–73.

42. Shin, J.H., et al., alphaB-crystallin suppresses oxidative stress-induced astrocyte apoptosis by inhibiting caspase-3 activation. Neurosci Res, 2009. 64(4): p. 355–61.

43. Kim, J.Y., et al., Alpha B-Crystallin Overexpression Protects Oligodendrocyte Precursor Cells Against Oxidative Stress-Induced Apoptosis Through the Akt Pathway. J Mol Neurosci, 2020. 70(5): p. 751–758.

44. Christopher, K.L., et al., Alpha-crystallin-mediated protection of lens cells against heat and oxidative stress-induced cell death. Biochim Biophys Acta, 2014. 1843(2): p. 309–15.

45. Smith, A.A., et al., Gene duplication and separation of functions in alphaB-crystallin from zebrafish (Danio rerio). FEBS J, 2006. 273(3): p. 481–90.

46. Koteiche, H.A., et al., Species-Specific Structural and Functional Divergence of alpha-Crystallins: Zebrafish alphaBa- and Rodent alphaA(ins)-Crystallin Encode Activated Chaperones. Biochemistry, 2015. 54(38): p. 5949–58.

47. Mukaigasa, K., et al., Genetic evidence of an evolutionarily conserved role for Nrf2 in the protection against oxidative stress. Mol Cell Biol, 2012. 32(21): p. 4455–61.

48. Mishra, S., et al., Loss of alphaB-crystallin function in zebrafish reveals critical roles in the development of the lens and stress resistance of the heart. J Biol Chem, 2018. 293(2): p. 740–753.

49. Zou, P., et al., A conserved role of alphaA-crystallin in the development of the zebrafish embryonic lens. Exp Eye Res, 2015. 138: p. 104–13.

50. Kramer, A., et al., Causal analysis approaches in Ingenuity Pathway Analysis. Bioinformatics, 2014. 30(4): p. 523–30.

51. Istvan, E.S. and J. Deisenhofer, Structural mechanism for statin inhibition of HMG-CoA reductase. Science, 2001. 292(5519): p. 1160–4.

52. Chen, Q.M. and A.J. Maltagliati, Nrf2 at the heart of oxidative stress and cardiac protection. Physiol Genomics, 2018. 50(2): p. 77–97.

53. Wang, J., et al., WEB-based GEne SeT AnaLysis Toolkit (WebGestalt): update 2013. Nucleic Acids Res, 2013. 41(Web Server issue): p. W77–83.

54. Yu, G., et al., DOSE: an R/Bioconductor package for disease ontology semantic and enrichment analysis. Bioinformatics, 2015. 31(4): p. 608–9.

55. Szklarczyk, D., et al., The STRING database in 2021: customizable protein-protein networks, and functional characterization of user-uploaded gene/measurement sets. Nucleic Acids Res, 2021. 49(D1): p. D605–D612.

56. Brown, S.L., et al., Activating transcription factor 3 is a novel repressor of the nuclear factor erythroid-derived 2-related factor 2 (Nrf2)-regulated stress pathway. Cancer Res, 2008. 68(2): p. 364–8.

57. Liu, P., et al., RPA1 binding to NRF2 switches ARE-dependent transcriptional activation to ARE-NRE-dependent repression. Proc Natl Acad Sci U S A, 2018. 115(44): p. E10352–E10361.

58. Widomska, J., et al., Cholesterol Bilayer Domains in the Eye Lens Health: A Review. Cell Biochem Biophys, 2017. 75(3-4): p. 387–398.

59. Widomska, J. and W.K. Subczynski, Why Is Very High Cholesterol Content Beneficial for the Eye Lens but Negative for Other Organs? Nutrients, 2019. 11(5).

60. Dotson, R.J., et al., Influence of Cholesterol on the Oxygen Permeability of Membranes: Insight from Atomistic Simulations. Biophys J, 2017. 112(11): p. 2336–2347.

61. Miyashita, T., et al., Evaluation of lens opacity due to inhibition of cholesterol biosynthesis using rat lens explant cultures. Toxicology, 2022. 465: p. 153064.

62. Kang, H., Z. Yang, and R. Zhou, Lanosterol Disrupts Aggregation of Human gammaD-Crystallin by Binding to the Hydrophobic Dimerization Interface. J Am Chem Soc, 2018. 140(27): p. 8479–8486.

63. Mori, M., et al., Lanosterol synthase mutations cause cholesterol deficiency-associated cataracts in the Shumiya cataract rat. J Clin Invest, 2006. 116(2): p. 395–404.

64. Wible, R.S., et al., Pharmacogenomics of Chemically Distinct Classes of Keap1-Nrf2 Activators Identify Common and Unique Gene, Protein, and Pathway Responses In Vivo. Mol Pharmacol, 2018. 93(4): p. 297–308.

65. Hybertson, B.M., B. Gao, and J.M. McCord, Effects of the Phytochemical Combination PB123 on Nrf2 Activation, Gene Expression, and the Cholesterol Pathway in HepG2 Cells. OBM Integr Compliment Med, 2022. 7(1).

66. Dimauro, I., et al., The role of alphaB-crystallin in skeletal and cardiac muscle tissues. Cell Stress Chaperones, 2018. 23(4): p. 491–505.

67. Rajasekaran, N.S., et al., Human alpha B-crystallin mutation causes oxido-reductive stress and protein aggregation cardiomyopathy in mice. Cell, 2007. 130(3): p. 427–39.

68. Yin, B., et al., CRYAB protects cardiomyocytes against heat stress by preventing caspase-mediated apoptosis and reducing F-actin aggregation. Cell Stress Chaperones, 2019. 24(1): p. 59–68.

69. Gupta, T. and M.C. Mullins, Dissection of organs from the adult zebrafish. J Vis Exp, 2010(37).

70. Duan, J., et al., Low-dose exposure of silica nanoparticles induces cardiac dysfunction via neutrophil-mediated inflammation and cardiac contraction in zebrafish embryos. Nanotoxicology, 2016. 10(5): p. 575–85.

71. Incardona, J.P. and N.L. Scholz, The influence of heart developmental anatomy on cardiotoxicity-based adverse outcome pathways in fish. Aquat Toxicol, 2016. 177: p. 515–25.

