## supporting information for "Interplay between Nrf2 and αB-crystallin in the lens and heart of zebrafish under proteostatic stress"

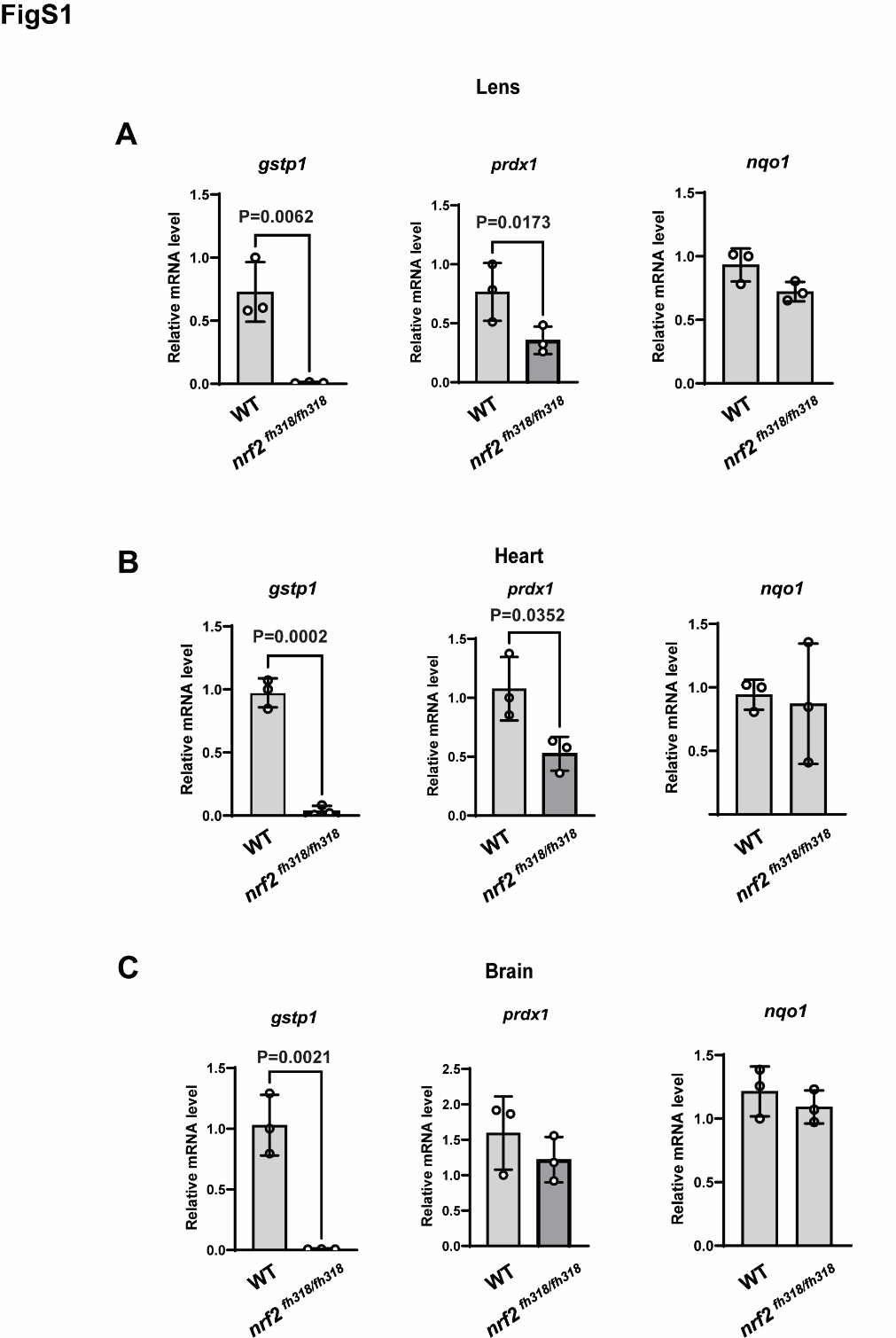

**Figure S1. Transcription changes of *nrf2* target genes in WT and nrf2^fh318/f318^.** The relative expressional changes of *gstp, prdx* and *nqo-1* between WT and nrf2^fh318/f318^ in lens **(A)**, heart **(B)**, and Brain **(C)** tissues were measured using qRT-PCR. Data are expressed as mean ± SD. *n*=3 for lens, heart, and brain tissues. Statistical significance was calculated using two-tailed t-test.

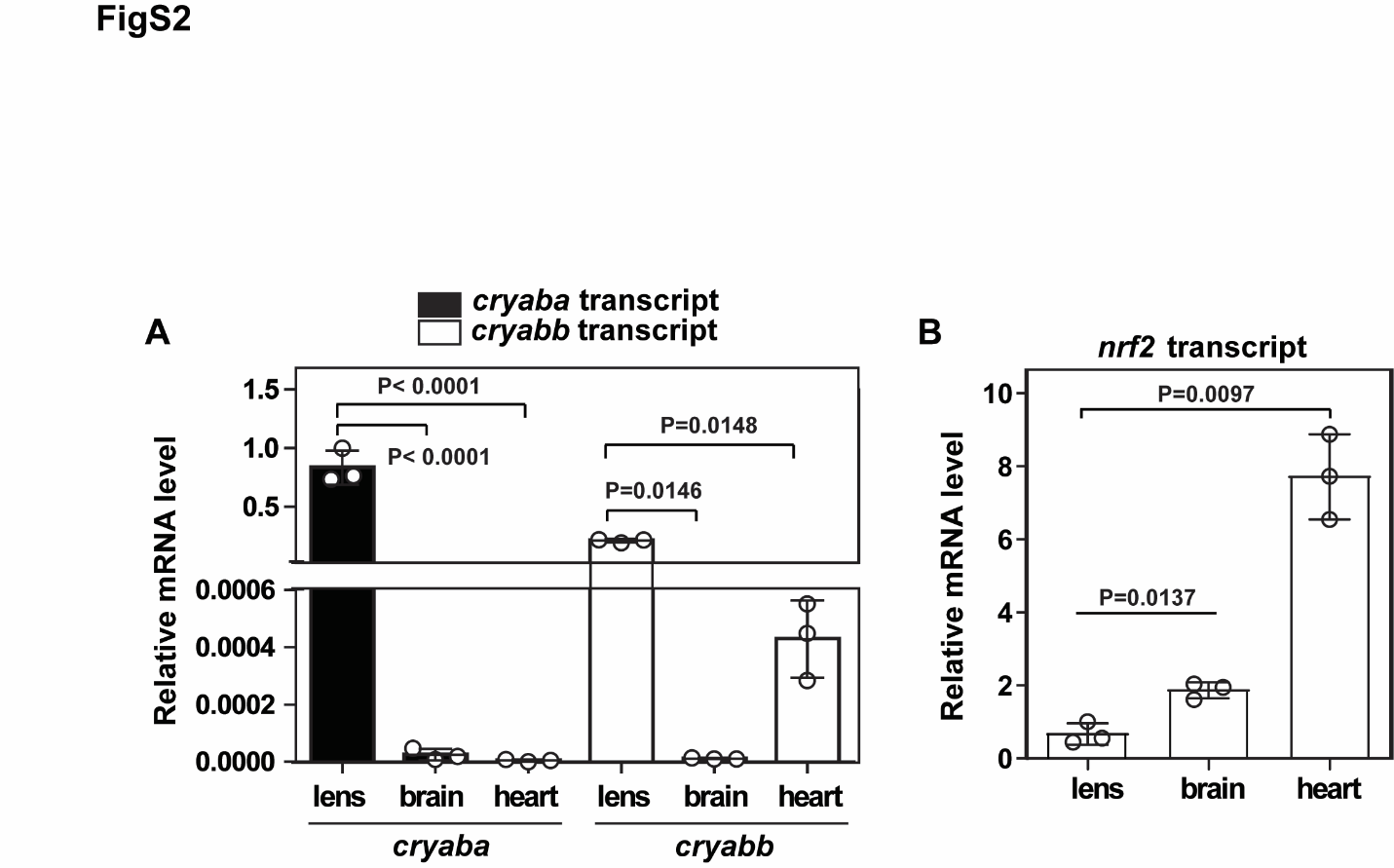

**Figure S2. Relative *cryaba, cryabb* and *nrf2* transcripts in different tissues.** The relative level of *cryaba*, *cryabb* (**A**) and *nrf2* (**B**) mRNA were compared using qRT-PCR analysis in different tissues. Statistical significance was calculated using ANOVA.

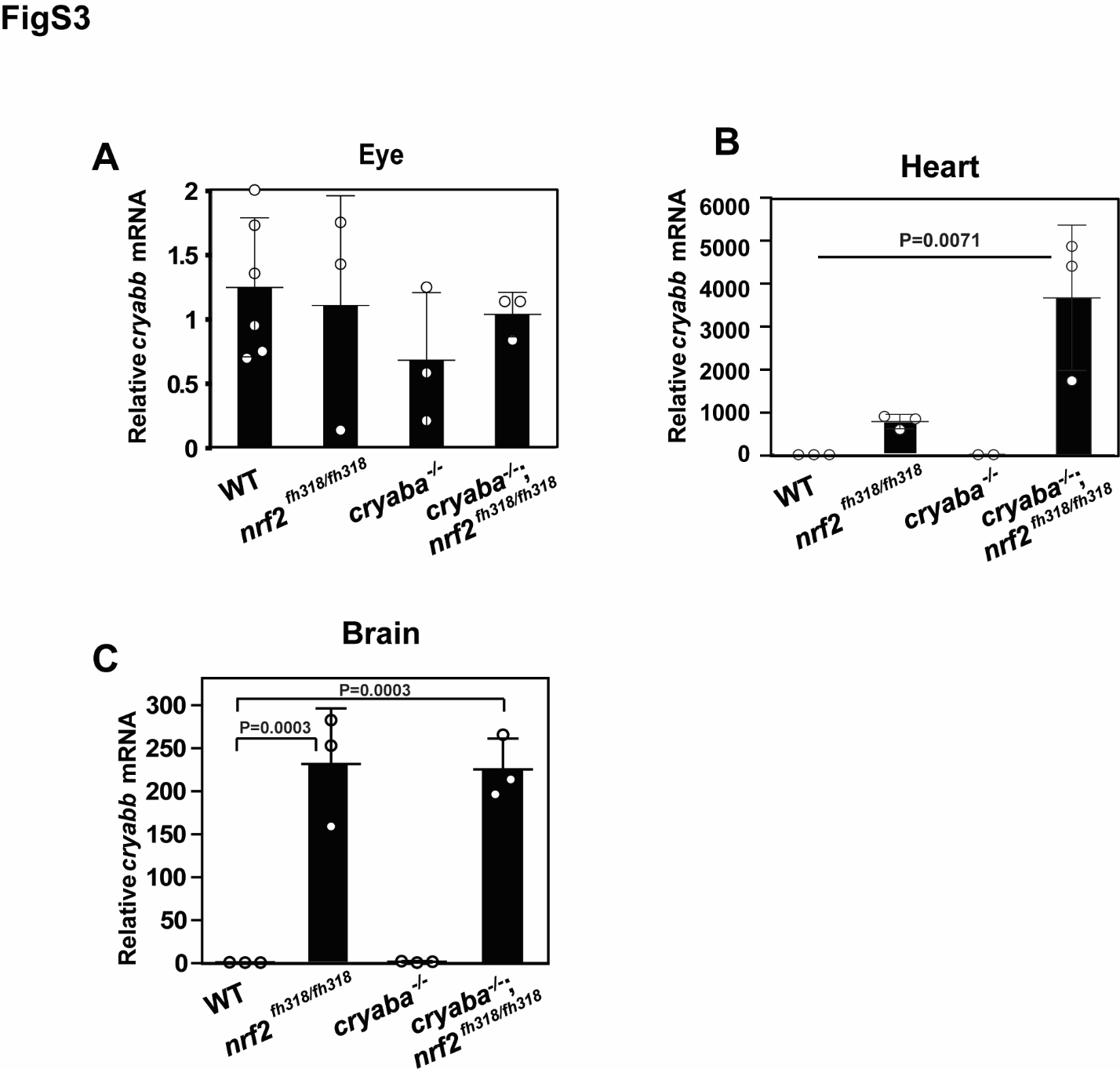

**Figure S3. Tissue-specific upregulation of *cryabb* in response to Nrf2 deficiency.** Relative mRNA expression of *cryabb* **(A)** in whole embryos at 4 dpf, **(B)** eyes, **(C)** heart, and **(D)** brain tissues of WT, *nrf2*^fh318/fh318^, *cryaba*^-/-^, and *cryaba*^-/-^;*nrf2*^fh318/fh318^. Data are expressed as mean ± SD. Statistical significance was calculated using one-way ANOVA.

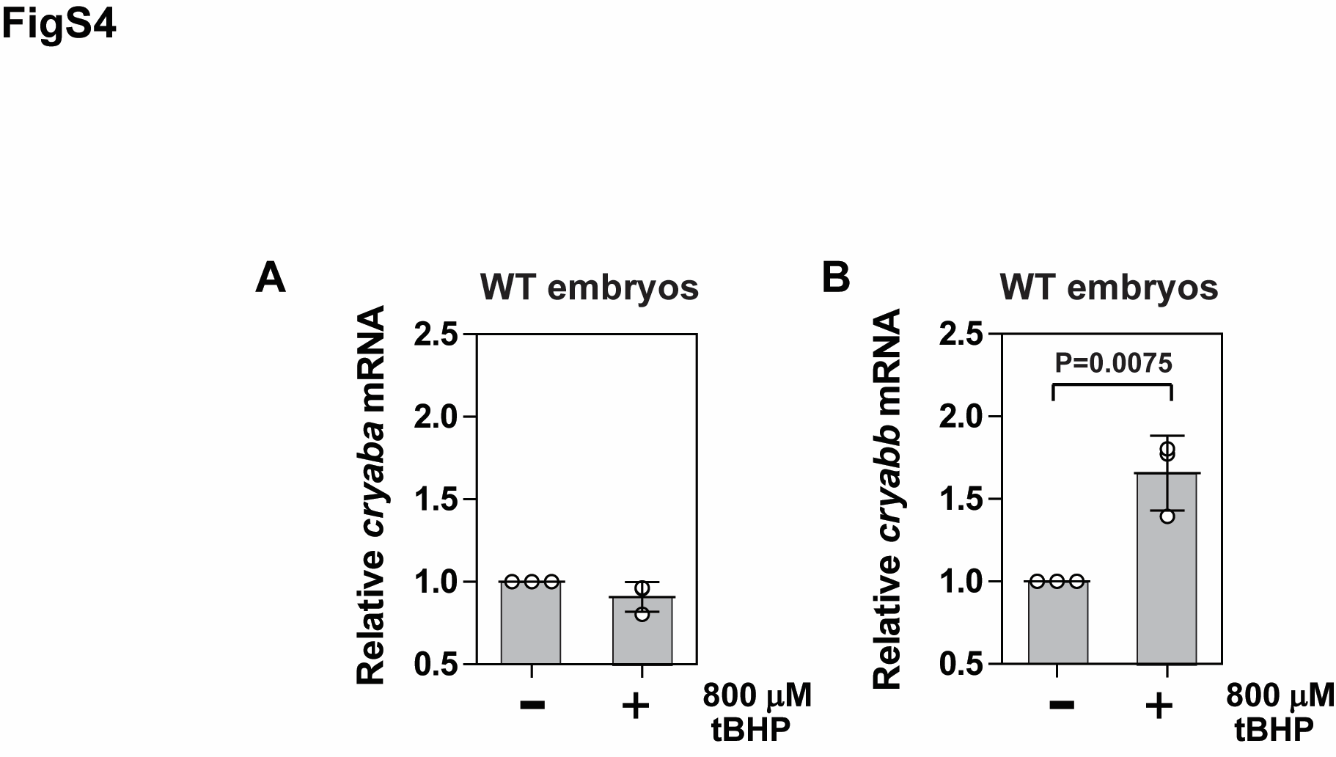

**Figure S4. Upregulation of *cryabb* transcript in response to tBHP treatment. (A)** Relative mRNA expression of *cryaba* and *cryabb* at 4 dpf were measured by qRT-PCR after treatment of 800 uM tBHP for two hours. Statistical significance was calculated using two-tailed t-test.

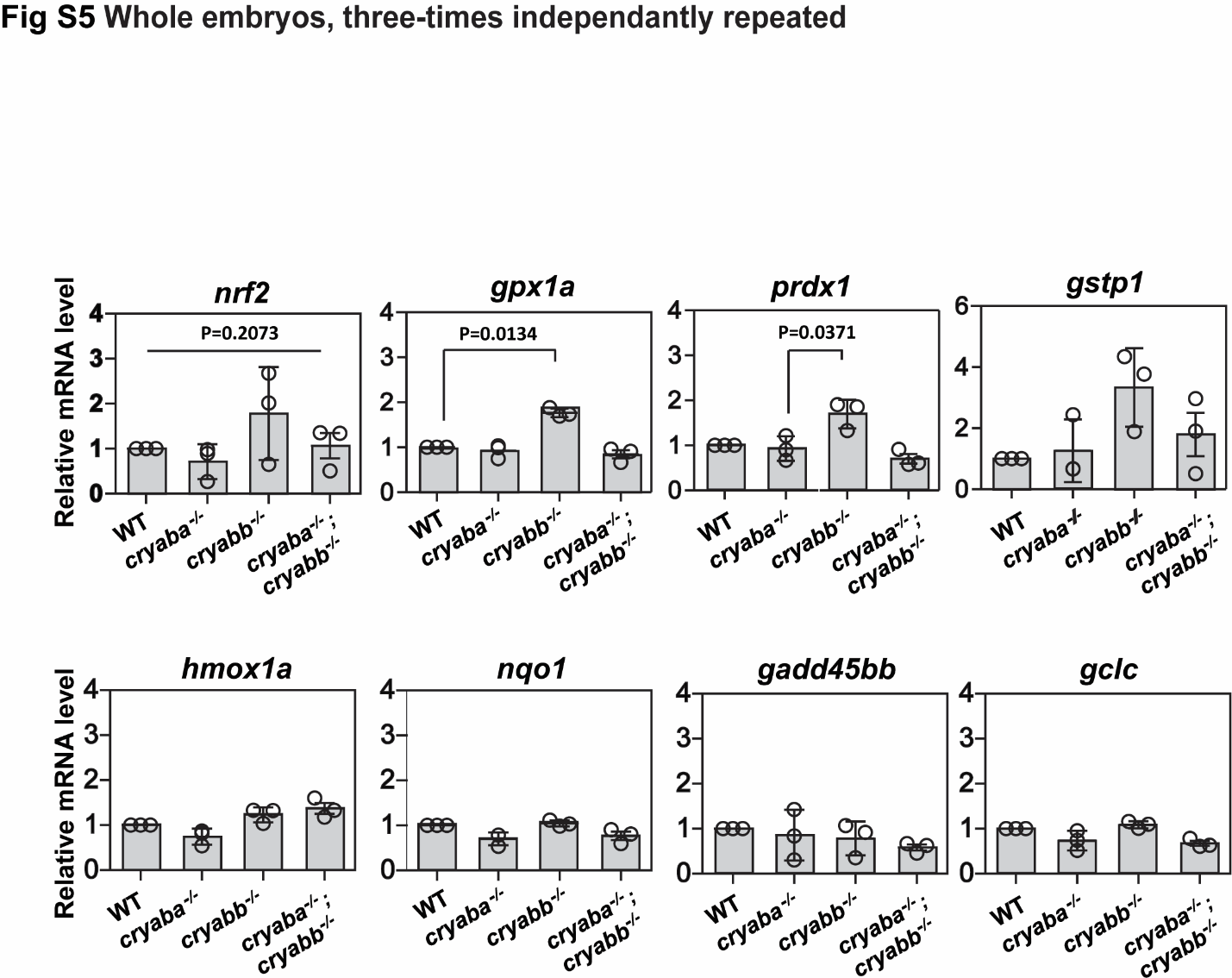

**Figure S5. Changes in the transcription of Nrf2 target genes in *cryab* mutants.** Relative levels of *nrf2, gpx1a, gstp1, prdx1, hmox1a, nqo1*, *gadd45bb*, and *gclc* in *cryaba^-/-^, cryabb^-/-^*, and *cryaba^-/-^;cryabb^-/-^* mutated embryos at 4 dpf were measured by qRT-PCR. Data are expressed as mean ± SD from 3 independent measurements. *P*-values were calculated using one-way ANOVA

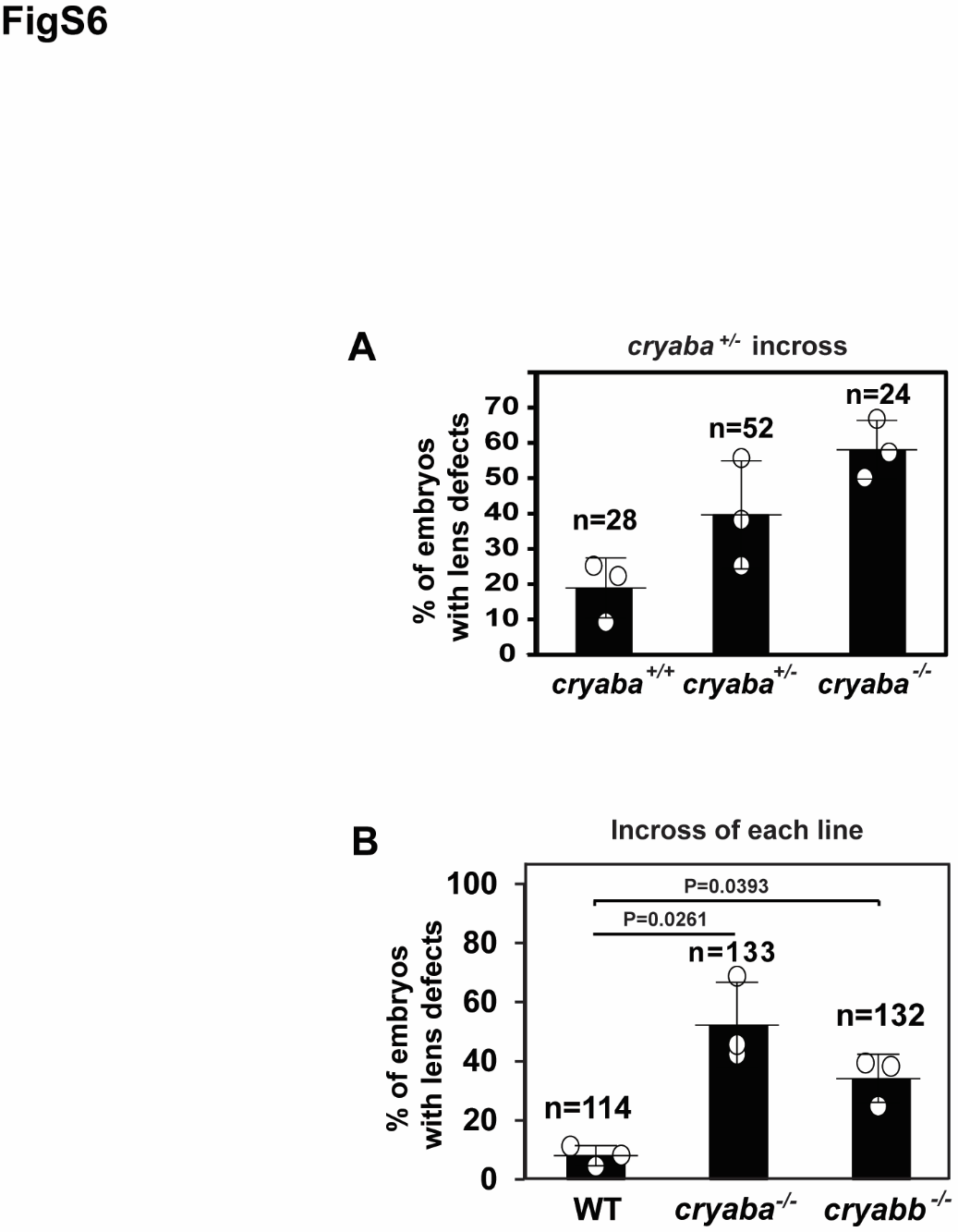

**Figure S6. Percentage of lens defect in *cryaba* and *cryabb* KO embryos. (A)** Lens of embryos from *cryaba*^+/-^ incross were screened at 4 dpf for lens abnormalities, and the genotype of *cryaba* was determined. **(B)** The percentage of lens defect at 4 dpf was compared among WT, *cryaba*^-/-^, and *cryabb*^-/-^. Statistical significance was calculated using one-way ANOVA.

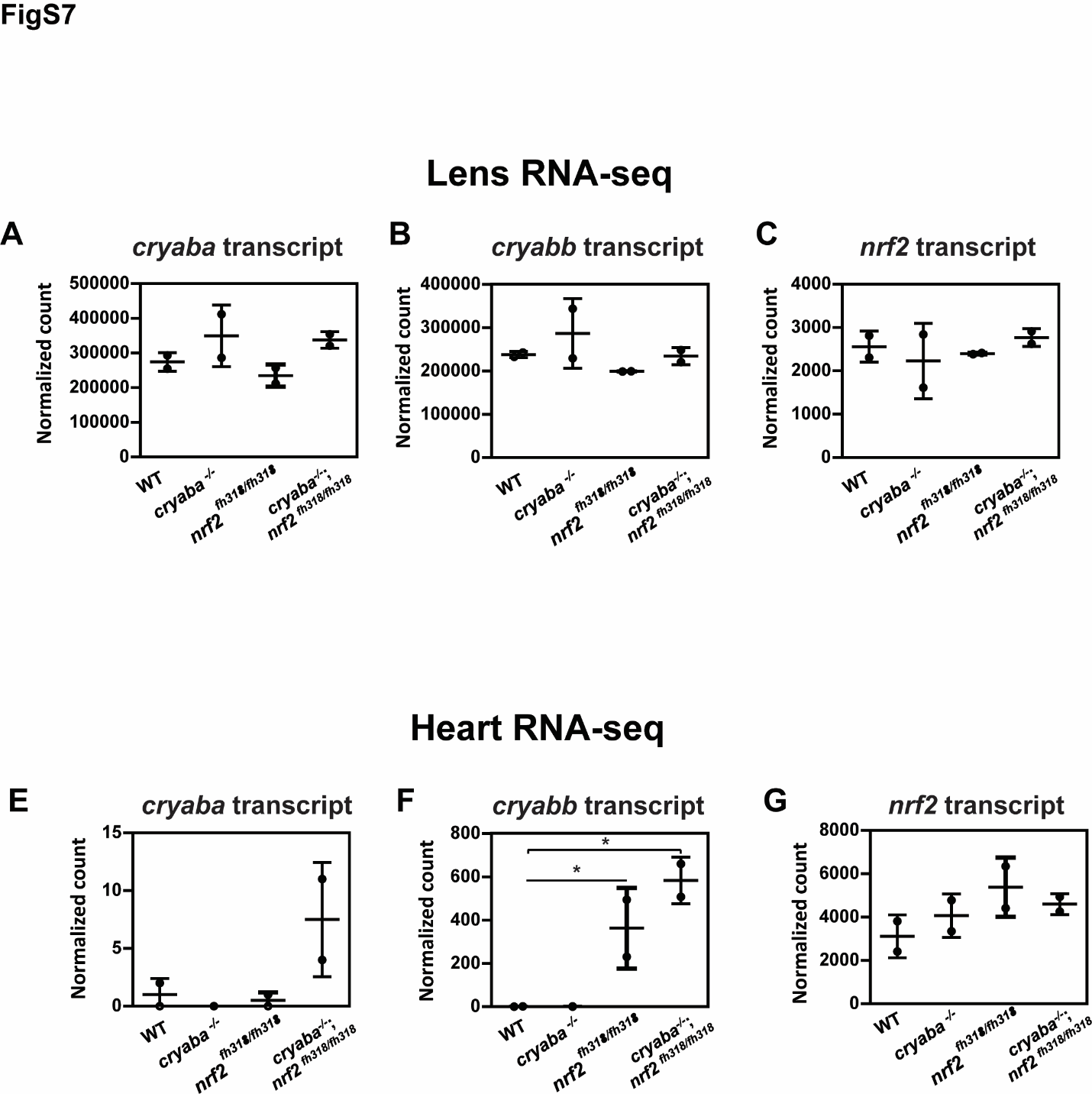

**Figure S7. The level of *cryaba*, *cryabb*, and *nrf2* mRNA in lens and heart tissues**. Normalized count values of *cryaba*, *cryabb*, and *nrf2* from lens RNA seq **(A-C)** and heart RNA-seq **(E-F)** are plotted as bar graphs. * Indicate False Discovery Rate (FDR) < 0.05.

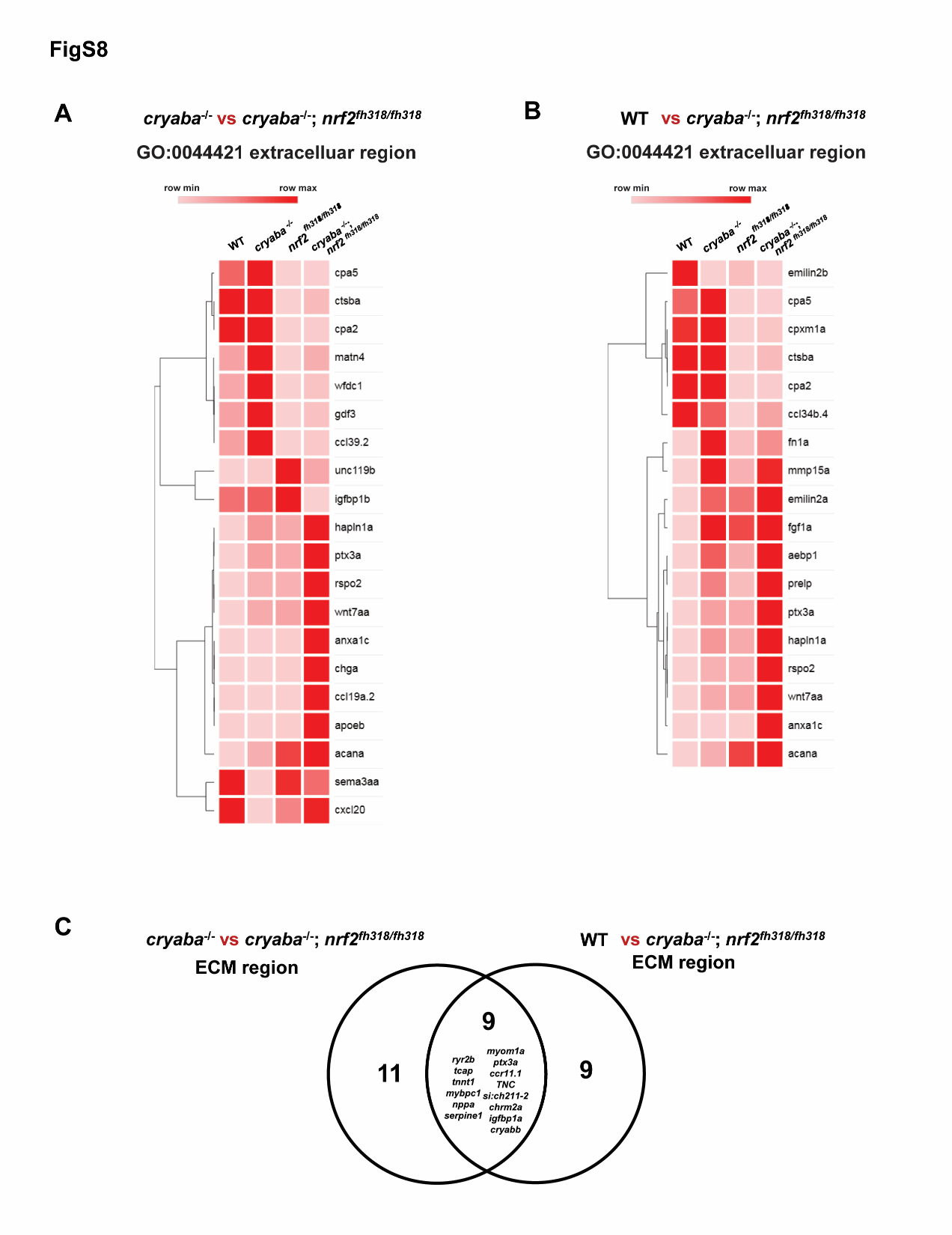

**Figure S8.** Heatmaps for the heart extracellular region GO cluster from *cryaba*^-/-^ versus *cryaba*^-/-^;*nrf2*^fh318/ fh318^ **(A)** and WT versus *cryaba*^-/-^;*nrf2*^fh318/ fh318^ **(B)**. **(C)** Venn-diagram analysis between the two GO clusters identifies significantly changed genes associated with ECM region in *cryaba*^-/-^;*nrf2*^fh318/ fh318^ compared with WT and *cryaba*^-/-^

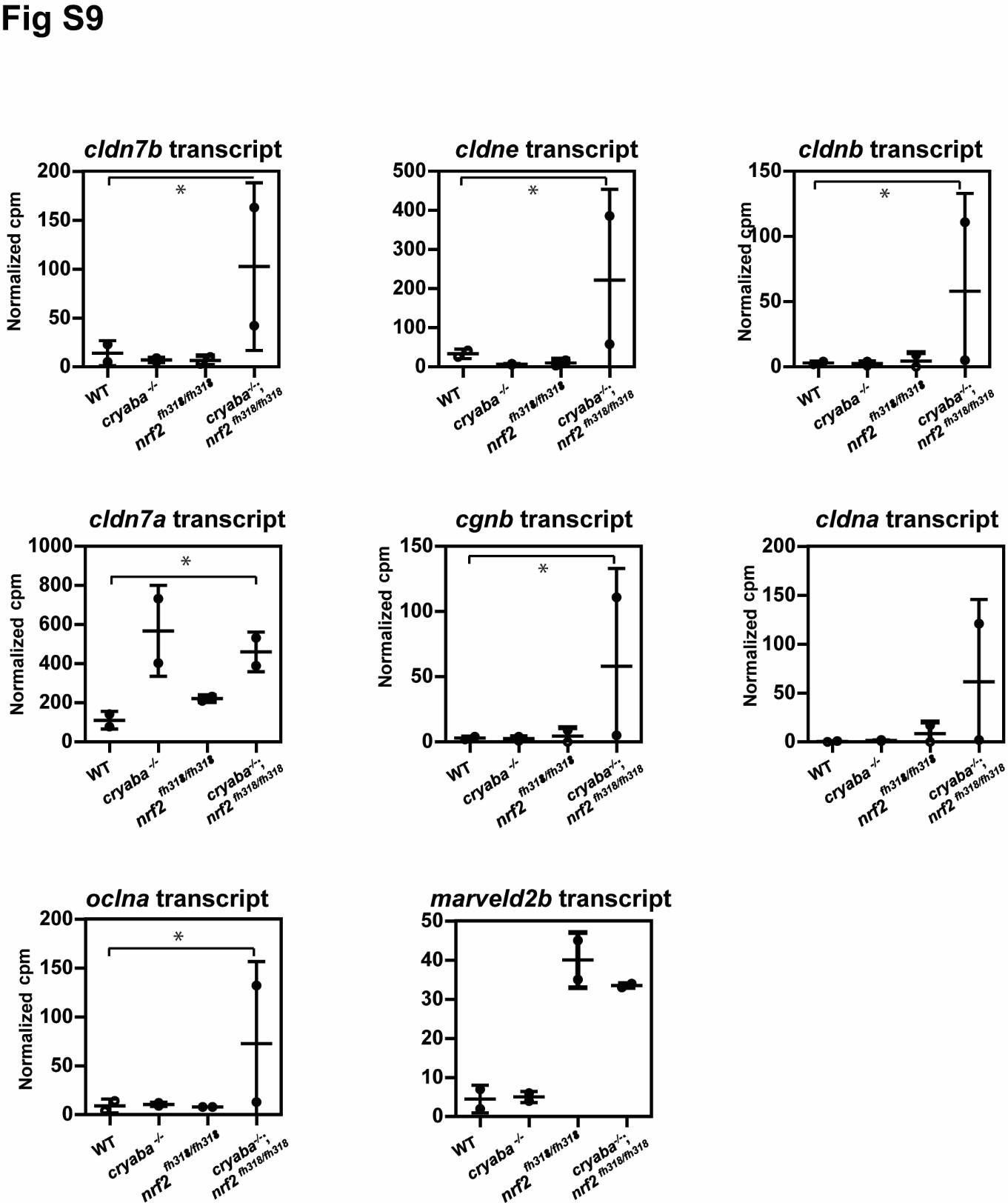

**Figure S9. Expression of genes in the tight junction GO cluster.** Normalized count values of each gene associated with tight junction are illustrated as bar charts from heart RNA seq data. * Indicate False Discovery Rate (FDR) < 0.05.

**Table S1. oligomer sequences for nrf2 pathway**

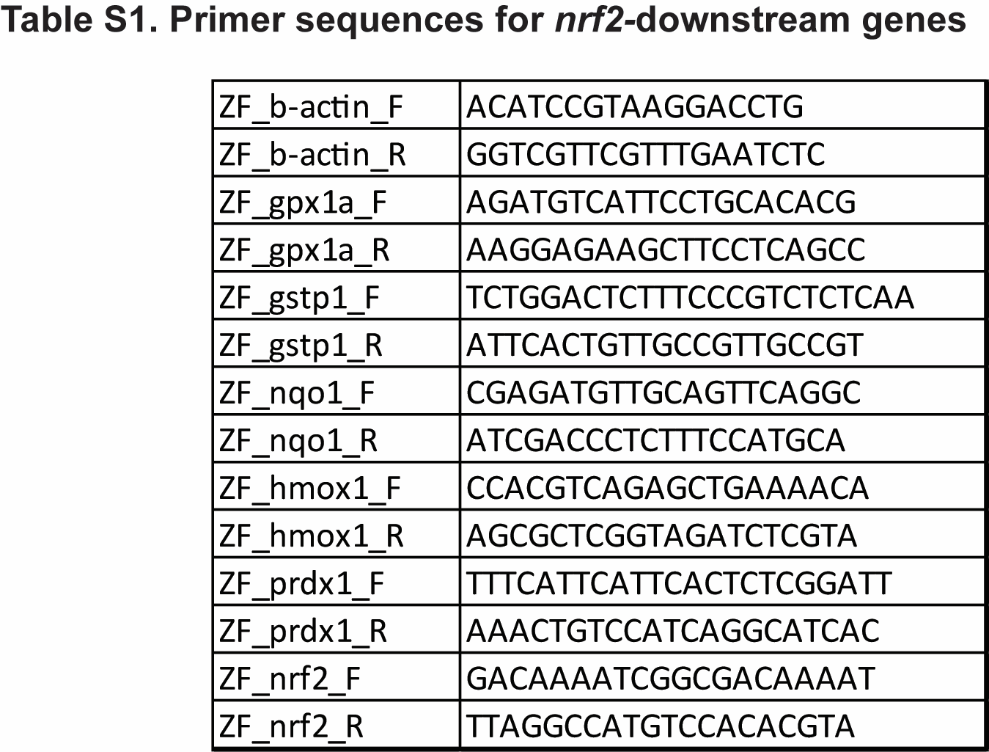

**Table S2. oligomer sequences for cholesterol biosynthesis pathway**

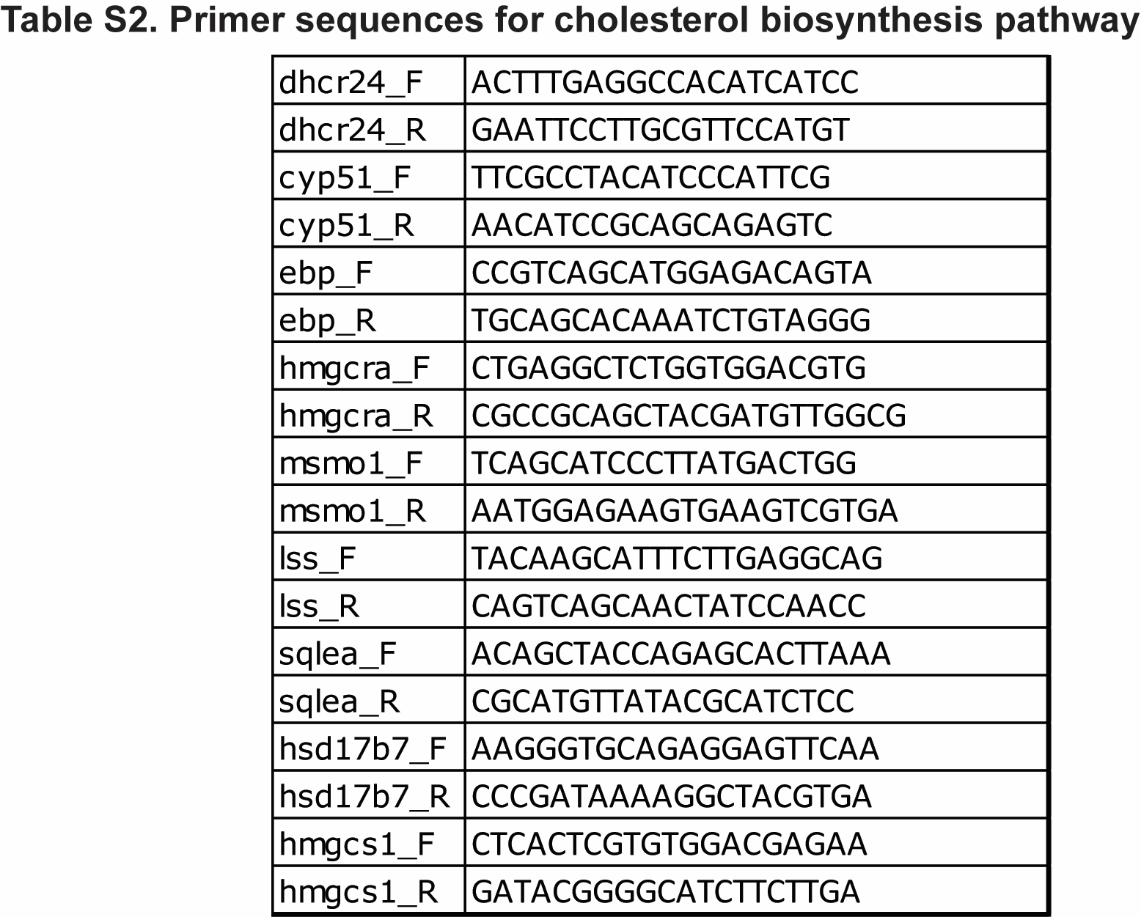

**Table S3. Normalized counts (cpm) of genes in extracelluar region part GO cluster from heart RNA-seq. Comparison between *cryaba*^-/-^ *versus* *cryaba*^-/-^;*nrf2*^fh318/ fh318^**

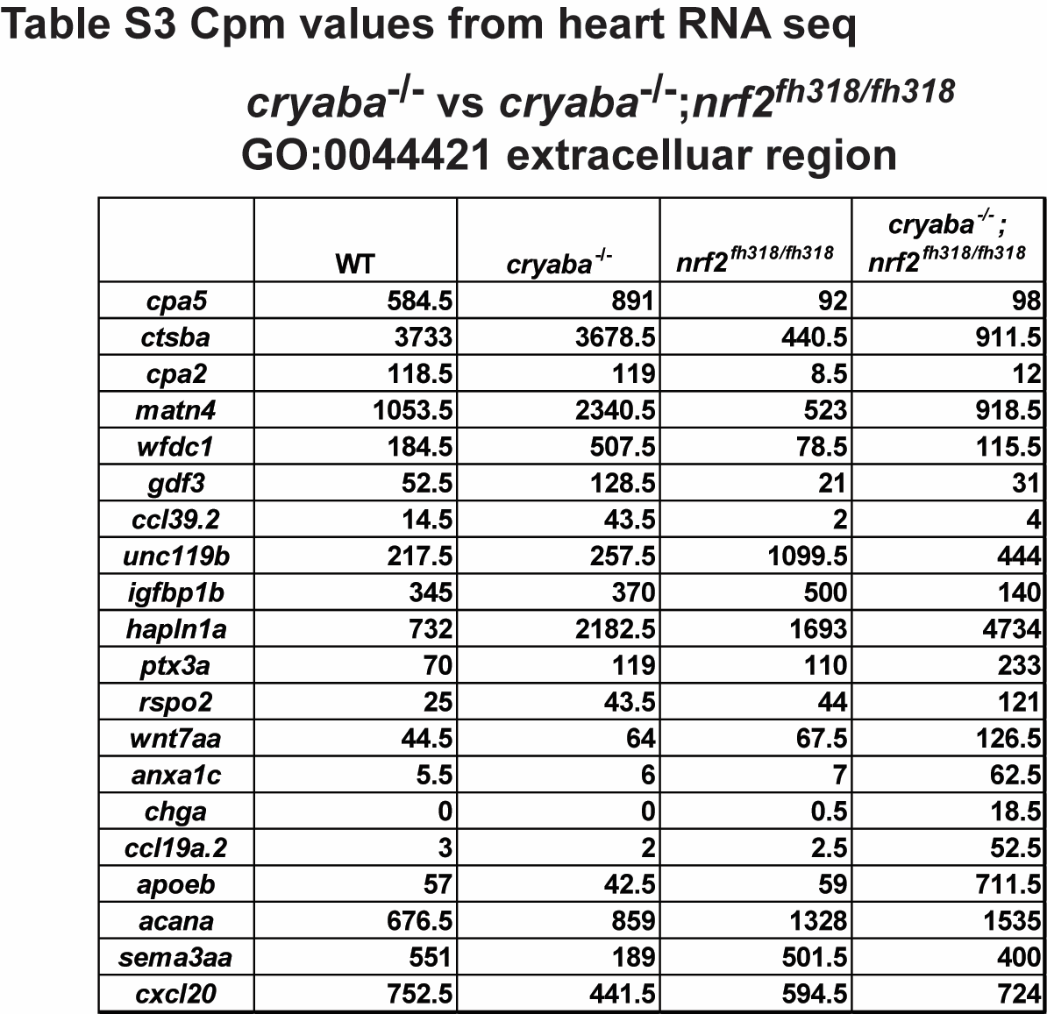

**Table S4. Normalized counts (cpm) of genes in the extracelluar region GO cluster from heart RNA-seq. Comparison between WT *versus* *cryaba*^-/-^;*nrf2*^fh318/ fh318^**

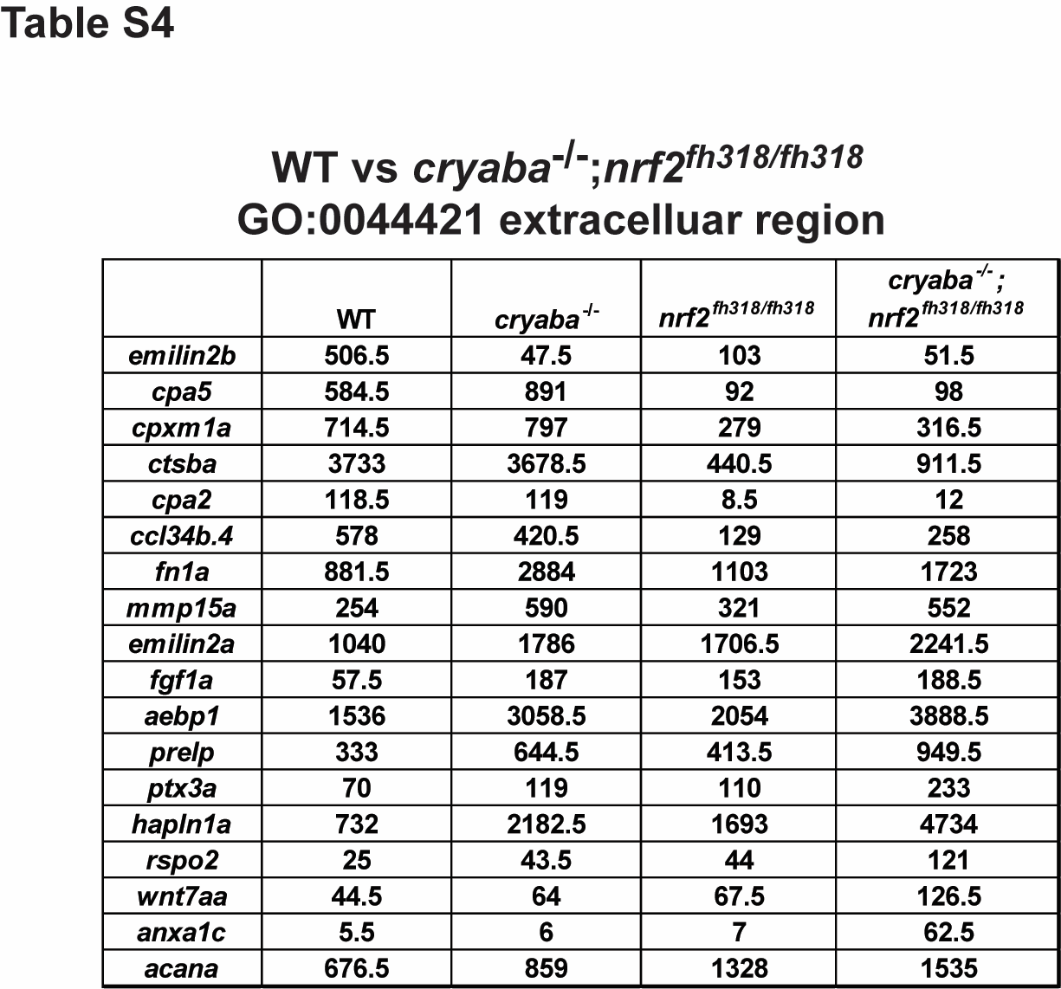
